# NLRP3 activators disrupt the endocytic AP2 complex and plasma membrane signaling

**DOI:** 10.64898/2026.02.24.707400

**Authors:** Stefan Ebner, Kshiti Meera Phulphagar, Yubell Alvarez, Lars Jürgenliemke, Fabian Frechen, Isabel Stötzel, Marta Lovotti, Matthew S. J. Mangan, Anil Akbal, Niels Schneberger, Annika Frauenstein, Tabea Klein, Jonathan J. Swietlik, Benedikt O. Gansen, Diana Fink, Andreas U. Lindner, Lea Nguyen, Isabelle Becher, Jonas Walter, Jessika Rollheiser, Anushka Kudaliyanage, Lukas Grätz, Oliver J. Gerken, Daniel Itzhak, Georg H. H. Borner, Maria C. Tanzer, Fraser Duthie, Rainer Stahl, Sebastian Kallabis, David Will, Mikhail M. Savitski, Agnes Schröder, Jonathan Jantsch, Gregor Hagelüken, Daniel F. Legler, Matthias Mann, Eicke Latz, Eva Kiermaier, Dagmar Wachten, Evi Kostenis, Felix Meissner

## Abstract

Organellar perturbations are linked to NLRP3 inflammasome activation, however, it remains unclear whether unrelated agonists converge on a common upstream pathway. Here, we traced intracellular organelle and protein movements by differential ultracentrifugation combined with mass spectrometry-based proteomics. We show that NLRP3 activators uniformly disrupt the endocytic Adaptor Protein 2 (AP2) complex, whereas other subcellular rearrangements are stimulus-specific. We discovered Dynasore as a K⁺-efflux–independent NLRP3 activator that engages this signaling node irrespective of endocytosis inhibition. Pharmacological and genetic perturbation of AP2 renders cells unresponsive to extracellular cues, blunting GPCR signaling, cAMP production, and chemotaxis, thereby enforcing a ’frozen’ signaling state that propagates NLRP3 inflammasome activation. Collectively, our study reveals a common surveillance checkpoint linking impaired plasma membrane signaling to the execution of inflammation and cell death.

**Graphical Abstract:** 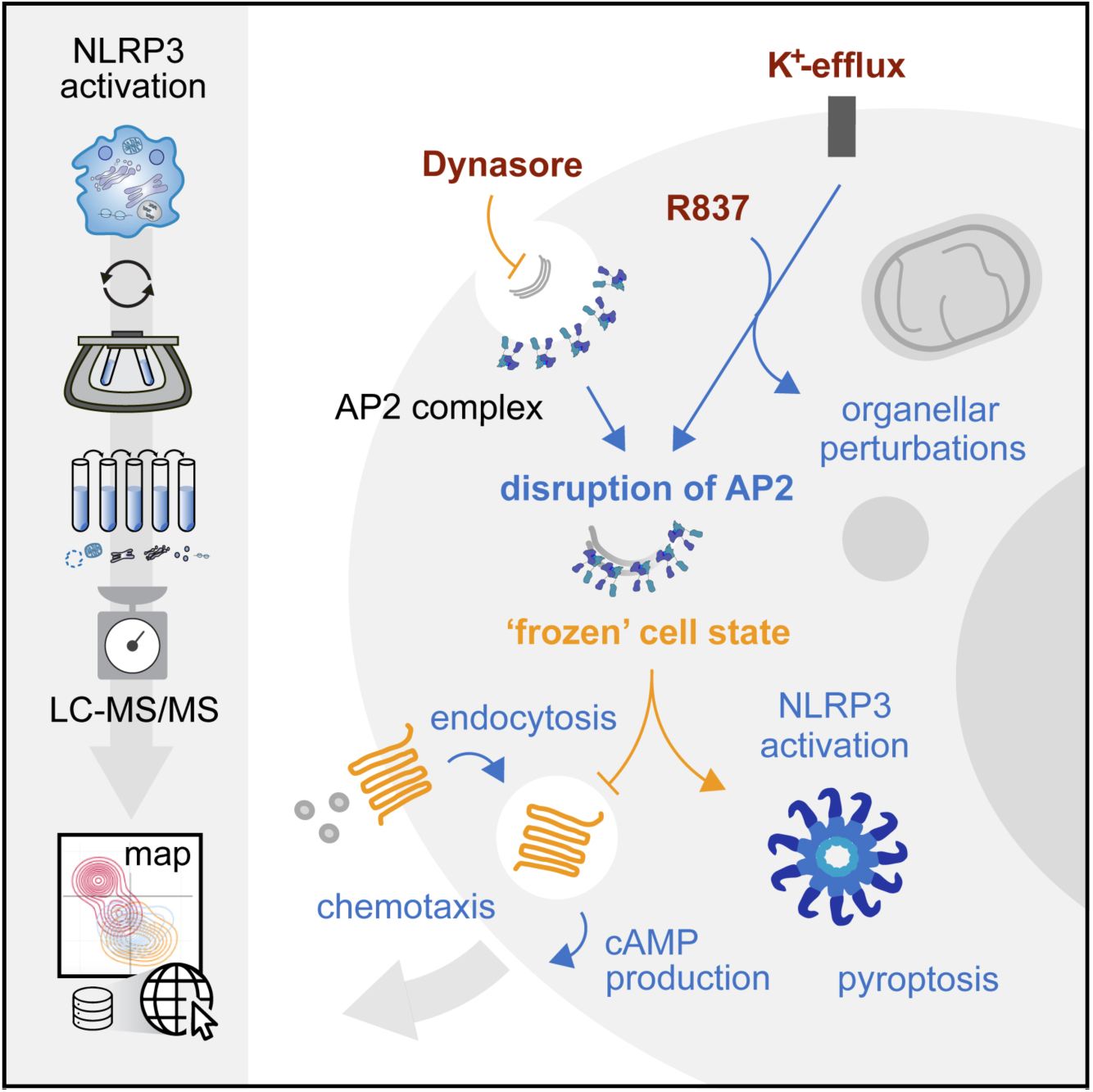

## Introduction

The NOD-like receptor family pyrin domain containing 3 (NLRP3) is a cytosolic innate immune sensor implicated in various inflammatory pathologies such as cryopyrin-associated periodic syndromes (CAPS) (*1*), type 2 diabetes, Alzheimer’s disease and atherosclerosis (*2*). Activation of NLRP3 triggers a highly inflammatory form of programmed cell death termed pyroptosis. This comprises the assembly of the adaptor protein ASC into specks, leading to caspase-1 activation and the subsequent pyroptotic release of IL-1β, IL-18, and other pro-inflammatory molecules through gasdermin D pores (*3–5*). As NLRP3 can be activated by structurally and mechanistically unrelated triggers, various indirect activation mechanisms have been proposed.

Cellular K^+^-efflux is a prominent danger signal shared between particulate matter, the ionophore Nigericin, ATP via P2X7R or KCNN4 downstream of the mechano-sensitive channel PIEZO1 (*6–8*). However, K^+^-efflux is dispensable for NLRP3 activation via imiquimod and the related compound CL-097 (*9*, *10*). Mitochondrial destabilization, which encompasses the suppression of ATP production, also provides a danger signal for activation and was proposed to disrupt the mitochondrial electron transport chain (*11*). Furthermore, dispersal of trans-Golgi network and endosomal trafficking can potentiate NLRP3 activation and provide sites for inflammasome formation after endosomal phospholipid accumulation chain (*11*). As a mechanism unifying these apparently distinct observations and a shared molecular event upstream of NLRP3 activation across agonists remained obscure, NLRP3 is often perceived as a broad sensor for organellar disruptions (*12*, *13*).

In this study, we set out to unbiasedly identify organellar perturbations required for NLRP3 activation. We traced spatiotemporal intracellular protein movements across NLRP3 agonists proteome-wide by dynamic organellar maps, combining subcellular fractionation with MS-based proteomics (*14*, *15*). To distinguish protein translocations upstream of inflammasome activation from those occurring during cell death, we employ cellular models deficient in pyroptosis execution. We find that all tested NLRP3 agonists uniformly disrupt the endocytic Adaptor Protein 2 (AP2) complex, while relocalizations of proteins from the mitochondria, the trans-Golgi network and endosomes are specific to few but not all stimuli. While pharmacologically interrogating clathrin-mediated endocytosis (CME), we serendipitously discovered Dynasore as a K^+^-efflux independent activator of NLRP3, which, in contrast to other CME inhibitors, also disrupts the AP2 complex. Using biotinylated Dynasore as a tool compound, we dissect its mode of action, identifying the nucleoside diphosphate kinases 1 and 2 (NME1, NME2) as molecular off-targets of Dynasore. We further delineate that NLRP3 agonists reduce responsiveness to external stimuli by interfering with GPCR internalization, cAMP-induced signal transduction and additionally chemotaxis, presumably through the disruption of the AP2 complex. Thus, our study reveals a conceptual paradigm in which innate immunity monitors the failure of essential plasma membrane signaling pathways.

## Results

### K^+^-efflux-dependent and -independent NLRP3 activation disrupts the AP2-complex upstream of pyroptosis

We set out to explore whether different NLRP3 activators induce shared organellar perturbations. We chose an experimental design where cell death is not executed thus permitting the dissection of common upstream events while preventing non-specific organellar disruptions and collateral damage downstream of inflammasome activation. To this end, we treated LPS-primed ASC or Caspase-1 deficient immortalized bone marrow-derived macrophages (iBMDMs) with the NLRP3 agonists Nigericin or imiquimod/ R837. We separated organelles by differential centrifugation into five fractions and analyzed them by mass spectrometry (MS)-based proteomics, as described previously (*14*, *15*) (Fig. 1A, S1A, S1B). Our workflow showed excellent reproducibility, evident by the separation of the fractions (Fig. 1B), and separation of organelles reflected by reference marker proteins (Fig. S1B). Upon NLRP3 activation, ASC displayed the most prominent change in subcellular localization, pelleting at lower centrifugation speeds, unbiasedly confirming speck formation by dynamic organellar maps. We identified 95 proteins, including ASC, consistently changing subcellular localization (Fig. S1C). Gene ontology (GO) enrichment analysis revealed NLRP3 activation-induced relocalizations of mitochondrial and trans-Golgi network (TGN) proteins (Fig. 1D), pelleting at higher centrifugation speeds and indicating a dispersal of the respective organelle in line with previous studies (*9*, *16*) (Fig. 1E). In addition, we detected even stronger shifts of clathrin/ AP2-complex proteins. AP2 acts as an obligate hetero-tetrameric protein complex which promotes clathrin cage-formation after recruitment to the plasma membrane during initiation of clathrin-mediated endocytosis (CME) (*17–20*) and had not been implicated in NLRP3 activation so far. While all complex subunits of AP2 reside in proximity to plasma membrane and endosomal proteins (Fig 1F), NLRP3 activation withdraws them from that localization (Fig. 1G, H), suggesting a deprival of functional AP2 from the plasma membrane.

**Figure 1.**
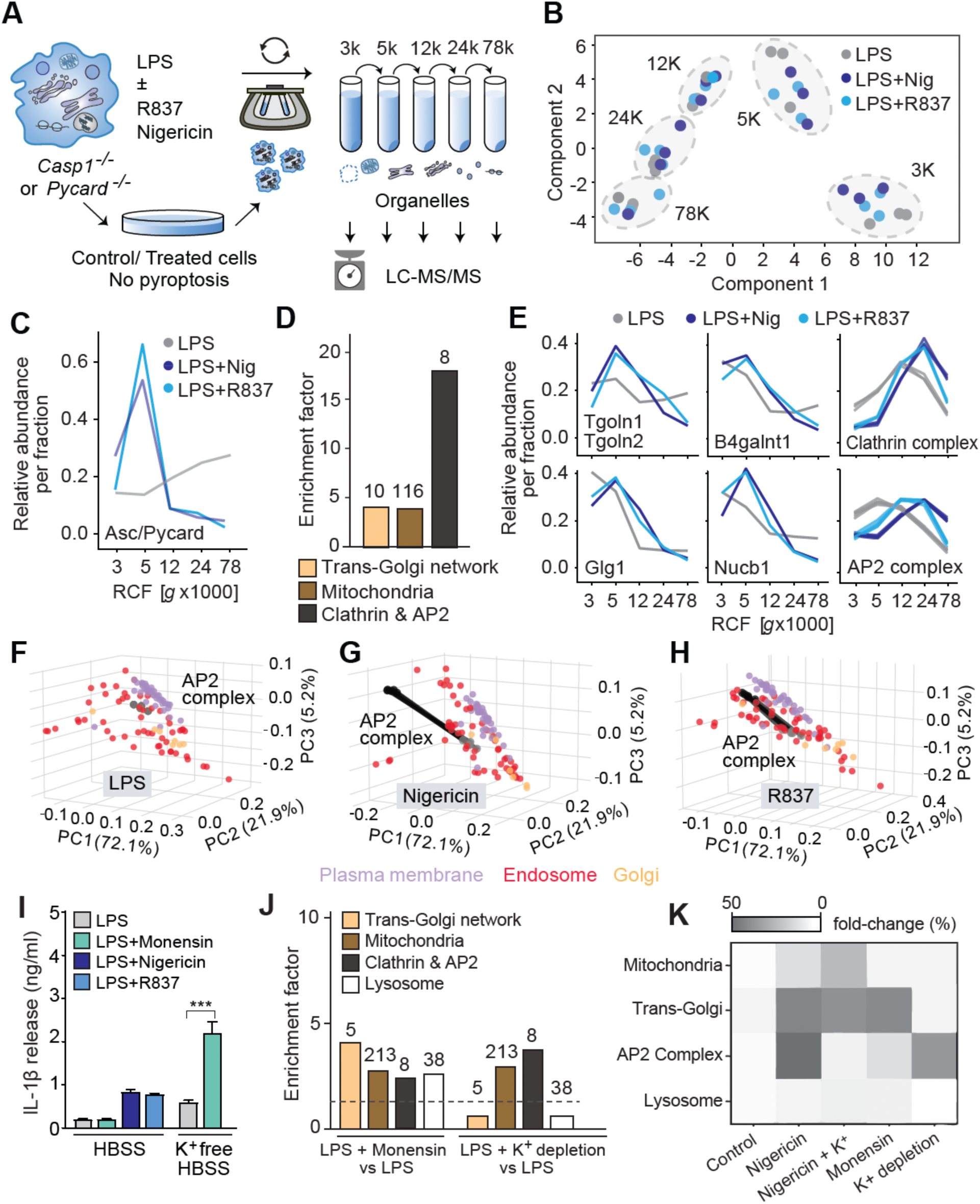
K^+^-efflux-dependent and -independent NLRP3 agonists disrupt the AP2-complex upstream of NLRP3. (A) Schematic illustration of the dynamic organellar maps workflow to dissect inflammasome activation upstream of NLRP3 in immortalized bone-marrow derived macrophages (iBMDMs) deficient in Caspase-1 or ASC. After activation and isotonic lysis, SILAC heavy labelled (Lys8, Arg10) iBMDMs are subjected to incremental ultracentrifugation, enriching different organellar fractions prior to addition of a light-labelled (Lys0, Arg0) whole-organellar reference spike-in. (B) Principal component analysis (PCA) of sub-organellar fractionation proteomes after TLR4-stimulation (priming, LPS 1 µg/ml, 4 h) or NLRP3 (LPS 4 h + Nigericin 2 µM/ R837 60 µg/ml for 120 min) stimulation in Caspase-1^-/-^ Balb/C iBMDMs. (C) Profile plot showing relative ASC distribution over sub-organellar fractions in Caspase-1^-/-^ iBMDMs. (D) Enrichment analysis for gene ontology cellular compartment (GOCC) terms and Uniprot keywords using Fisher exact test on proteins significantly changing (p < 0.05; FDR < 0.02) localization upon NLRP3 activation in ASC^-/-^ or Caspase-1^-/-^ iBMDMs. (E) Profile plots showing relative distribution of trans-Golgi-network proteins Tgoln1; Tgoln2, B4galnt1, Glg1 and Nucb1, as well as all clathrin or AP2 complex related proteins across organellar fractions from TLR4- or TLR4 and NLRP3-activated ASC^-/-^ iBMDM. (F) 3D-Principal-component-analysis of median normalized intensities per fraction showing position of median AP2-complex subunits (grey) in LPS-primed control with annotations for endosomal (red), Golgi- (orange) or plasma membrane proteins (blue). (G) 3D-Principal-component-analysis of median normalized intensities per fraction showing relocalization of AP2-complex subunits (grey-LPS, black-Nigericin, movement indicated by lines) upon 2 µM Nigericin treatment with annotations for endosomal (red), Golgi- (orange) or plasma membrane proteins (violet). (H) As in (H) for 60 µg/ml R837. (I) IL-1β release of wt C57Bl6 BMDMs in Hank’s balanced salt solution (HBSS) (5 mM KCl) or K^+^-free HBSS (0 mM KCl, + 5 mM NaCl) ± 2 µM monensin. Statistics indicate significance by student’s two-tailed t-test (*** = P < 0.001). (J) Enrichment analysis of GOCC terms and Uniprot keywords using a Fisher exact test on all proteins significantly (p < 0.05; FDR < 0.02) changing localization upon monensin treatment or exposure to K^+^-free HBSS in ASC^-/-^ or Caspase-1^-/-^ iBMDMs. (K) Heatmap showing combined absolute fold-changes of organellar marker proteins belonging to mitochondria, TGN, AP2 or lysosomes from dynamic organellar maps in ASC-deficient BMDMs treated with LPS (control), 2 µM Nigericin, 2 µM Nigericin + 80 µM KCl, 2 µM monensin or 2 µM monensin in K^+^-free buffer.

To examine whether this displacement is NLRP3 specific and not a consequence of broad organellar perturbations, we compared the K^+^-ionophore Nigericin and the supposedly K^+^-independent agonist R837 to the Na^+^-ionophore monensin, which induces pH perturbations similar to Nigericin without activating NLRP3 (*21*), or to a K^+^-free buffer, which triggers NLRP3 activation solely via osmotic gradients (*22*). K^+^-depletion was sufficient to trigger IL-1β release from iBMDMs while monensin boosted NLRP3 activation in combination with K^+^-depletion (Fig. 1I). Dynamic organellar maps in ASC deficient iBMDMs showed that monensin triggered broad organellar perturbances, affecting mitochondria, ribosomes, lysosomes and also the TGN (Fig. 1J, S1E) while in contrast, K^+^-depletion specifically redistributed AP2-complex/ clathrin family proteins and, to a lesser extent, mitochondrial proteins (Fig 1J, K, S1D). Further, preventing ion fluxes by a surplus of extracellular KCl during Nigericin-induced NLRP3 activation markedly reduced AP2 relocalization, while TGN and mitochondrial perturbations persisted (Fig. 1K). Together our results point to specific spatiotemporal redistribution of AP2 across NLRP3 agonists. Our spatial proteomics data are publicly accessible through an interactive website https://immprot.org/nlrp3mapper.

### Dynasore is a K^+^-efflux independent NLRP3 agonist that triggers AP-2 relocalization from the plasma membrane

Depletion of intracellular K^+^ prevents the formation of clathrin-coated pits and results in the generation of AP2-rich vesicles, which detach from the plasma membrane (*23*). We explored whether this arrest of pit formation or rather a general disruption of CME triggers NLRP3 activation and the relocalization of AP2. We exposed BMDMs to Dynasore *(N*-[(*E*)-(3,4-dihydroxyphenyl)methylideneamino]-3-hydroxynaphthalene-2-carboxamide) or the derivative Dyngo-4a (3-hydroxy-*N*-[(*E*)-(2,4,5-trihydroxyphenyl)methylideneamino]naphthalene-2-carboxamide), both of which inhibit Dynamin GTPase activity and prevent endocytosis (*24–27*). Strikingly, Dynasore induced pyroptosis in a dose-dependent manner to comparable levels as Nigericin and extracellular ATP, while Dyngo-4a did not (Fig. 2A). This also held true for human monocyte-derived macrophages (Fig S2A). Dynasore and Dyngo-4a differ only in one hydroxyl group, and while both block endocytosis, this structural difference appears responsible for NLRP3 activation by Dynasore.

**Figure 2.**
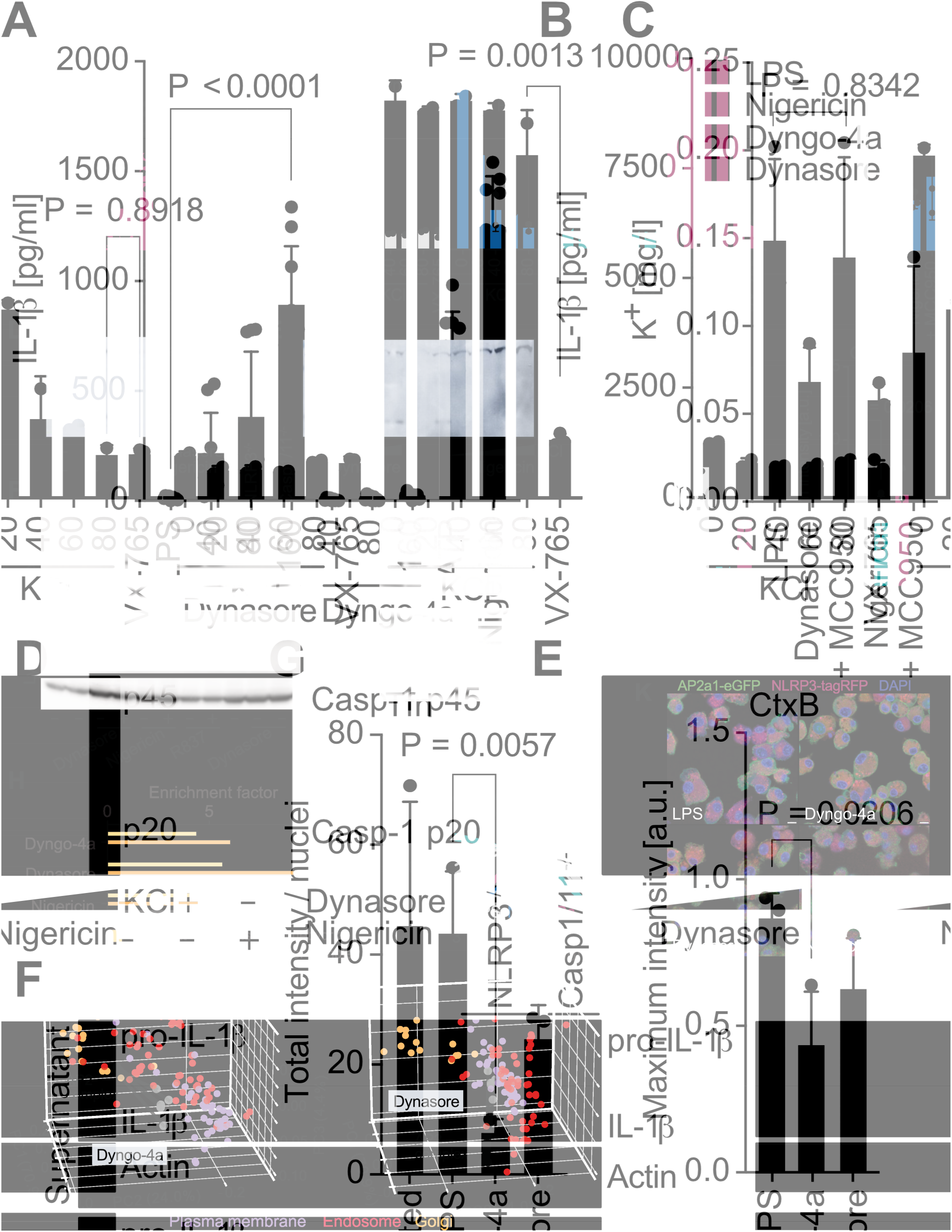
Dynasore is a K^+^-efflux independent NLRP3 agonist that triggers AP-2 relocalization from the plasma membrane. (A) IL-1β release of LPS-primed *wt* Balb/c iBMDMs treated with Dyngo-4a, Dynasore (40/ 80/ 160 µM), ATP (2 mM) or Nigericin (2 µM) for 90 min; n = 3. (B) IL-1β release of LPS-primed C57Bl6 BMDMs stimulated with Nigericin (2 µM), Dyngo-4a (80 µM) or Dynasore (160 µM) in presence of excessive KCl (0-80 mM) or the Caspase-1 inhibitor VX-765 (10 µM); n = 3 (C) Quantification of intracellular K^+^ via ion-selective electrode measurement. MCC-950 indicates Nlrp3 inhibitor treatment. (D) Representative Western blot for cleaved (p20) and uncleaved (p45) caspase-1 from primary C57Bl6 BMDM supernatants; n = 3 (E) As in C, but in presence of increasing concentrations of KCl; n = 3. (F) Representative Western blot for mature and pro-IL-1β in presence or absence of excess 80 mM KCl in *wt* Balb/C iBMDMs, NLRP3- or Caspase-1/11 deficient iBMDMs, treated with 80 µM Dynasore, 10 µM Nigericin, 60 µg/ml R837 (left) or 20/ 40/ 80 µM Dynasore or Dyngo-4a (right); n = 2. (G) Relative AlexaFlour-647 labelled transferrin (Tfn, left) or AlexaFlour-594 labelled Choleratoxin-B (CtxB, right) uptake in ASC^-/-^ iBMDMs in the presence of Dyngo-4a or Dynasore; n = 3 (H) Enrichment analysis for GOCC terms and Uniprot keywords using Fisher exact test on proteins significantly (p < 0.05; FDR < 0.02) changing localization in ASC^-/-^ iBMDMs in presence of Dyngo-4a (80 µM), Dynasore (160 µM) or Nigericin (2 µM) relative to LPS-primed control. (I) Profile plots showing relative AP2-complex subunit (AP2a1, AP2a2, AP2b1, AP2m1, AP2s1) distribution across fractions. (J) 3D-Principal-component-analysis showing transition of AP2-complex subunits (grey-LPS, black-Nigericin, movement indicated by lines) upon 80 µM Dyngo-4a (left) or 160 µM Dynasore (right) treatment with annotations for endosomal (red), Golgi- (orange) or plasma membrane proteins (violet). (K) Representative confocal microscopy images of LPS-primed ASC^-/-^ iBMDMs stably expressing functional AP2a1-eGFP-AP2b1 fusion protein and transiently expressing NLRP3-tagRFP, co-stained with DAPI. Statistics indicate significance by student’s two-tailed t-test (A, B, C, F, G).

We next excluded that Dynasore activates NLRP3 by permeabilizing the plasma membrane and allowing subsequent K^+^-efflux, as suggested (*28*). Excessive KCl in the medium did not inhibit Dynasore-induced NLRP3 activation (Fig. 2B). Likewise, intracellular K^+^-levels did not change by Dynasore treatment, and only decreased when pyroptosis was executed (Fig. 2C). Furthermore, Dynasore treatment resulted in similar caspase-1 and IL-1β cleavage as compared to Nigericin in primary BMDMs (Fig. 2D, F), which, unlike Nigericin, persisted despite addition of excessive KCl (Fig. 2E, F). Dynasore activates NLRP3 specifically, reflected in abrogated IL-1β release upon Caspase-1 inhibitor (VX-765) (Fig. 2B), NLRP3-inhibitor (MCC-950) treatment (Fig. S2A) or in NLRP3-deficient iBMDMs (Fig. 2F, S2H), but not NLRP1 or AIM2 knocked down iBMDMs (Fig. S2I). Dynasore also did not affect the secretion of conventionally released cytokines such as IL-6 or TNF (Fig. S2F). Together, this establishes Dynasore as a bona fide K^+^-efflux-independent NLRP3 activator.

Dynasore reportedly has off-target effects compared to its derivate Dyngo-4a. We reasoned that it may engage additional cellular targets that mediate NLRP3 activation, presumably independent of endocytosis inhibition (*25–28*). Indeed, both compounds prevented CME and clathrin-independent endocytosis to comparable levels, as quantified by the uptake of transferrin or Cholera toxin B (Fig. 2G), which are endocytosed clathrin-dependently or - independently, respectively. To evaluate differences in perturbed cellular pathways, we compared either compound to Nigericin by dynamic organellar maps in ASC-deficient iBMDMs. GO-term enrichment analysis confirmed the relocalizations of trans-Golgi proteins, proteins involved in CME (“Coatedpit”) and, intriguingly, AP2 complex members only upon Dynasore and Nigericin treatment (Fig. 2H). Nigericin and Dynasore indeed displayed very comparable relocalization profiles for AP2 (Fig. 2I, S2B), displacing all members from both the plasma membrane and the endosome (Fig. 2J), while Dyngo-4a repositioned them towards the endosome. Only 10 proteins consistently changed their localization upon both Dynasore and Nigericin treatment that were not shared with Dyngo-4a (Fig. S2G). These comprise several subunits of AP2 and EPS15, the latter acting in unison with AP2 during CME. Nigericin and R837 also relocalized AP2 and associated proteins like EPS15, as well as clathrins, GAK and Picalm, pointing to a central signaling nexus upstream of inflammasome activation (Fig. S1C). To rule out that AP2 relocalization is triggered by Dynasore’s known side effect on endolysosomal acidification, we excluded that the relocalization of AP2 is triggered by chloroquine, bafilomycin A1, the proton pump uncoupler FCCP or the PI3-kinase and macropinocytosis inhibitor Wortmannin. (Fig. S2C). We further excluded ROS production and mitochondrial Dynamin DRP1 as mediators of Dynasore-induced NLRP3 activation (Fig. S2D, E). Collectively, our data show that Dynasore, but not Dyngo-4a displaces the AP2 complex from the plasma membrane - a specific perturbation shared among NLRP3 agonists.

### Dynasore interacts with NME1/2 and - like other NLRP3 agonists - activates the NLRP3 inflammasome AP2-complex-dependently

We wondered whether AP2 directly engages with NLRP3 after relocalization. To test this, we introduced an internally tagged functional eGFP-AP2-fusion construct (*29*) into iBMDMs together with an RFP-tagged NLRP3 fusion and examined whether colocalization occurs. We recapitulated the Dynasore-induced movement of AP2 away from the plasma membrane, but observed no colocalization of AP2 and NLRP3 (Fig. 2K), arguing against a direct activation mechanism driven by interaction. Conversely, we investigated whether Dynasore directly interacts with components of the AP2 complex. Since the movement of AP2 also occurs for R837 and Nigericin, we surmised that the identification of Dynasore’s molecular target upstream of inflammasome activation might provide valuable entry points in distinguishing a common upstream protein target which mediates the relocalization of AP2. To this end, we synthesized analogs of Dynasore and Dyngo-4a containing a biotin affinity handle on the opposite side of the functionally distinct groups (Fig. S3A), for proteomics affinity purification experiments. We identified three class I nucleoside diphosphate kinases specifically NME1, NME2 and to lesser extent NME4 among the top10-differentially enriched proteins for biotin-Dynasore in LPS-primed BMDMs, which were not bound by biotin-Dyngo-4a (Fig. 3A, B). Congruently, 1D-annotation enrichment analysis shows strong enrichment of proteins involved in purine- or nucleotide metabolism, to which NMEs belong (Fig. S3B). Binding predictions using AlphaFold3 locate Dynasore in a pocket required for NME1’s kinase activity with an ipTM score of 0.75 (Fig 3D).

**Figure 3.**
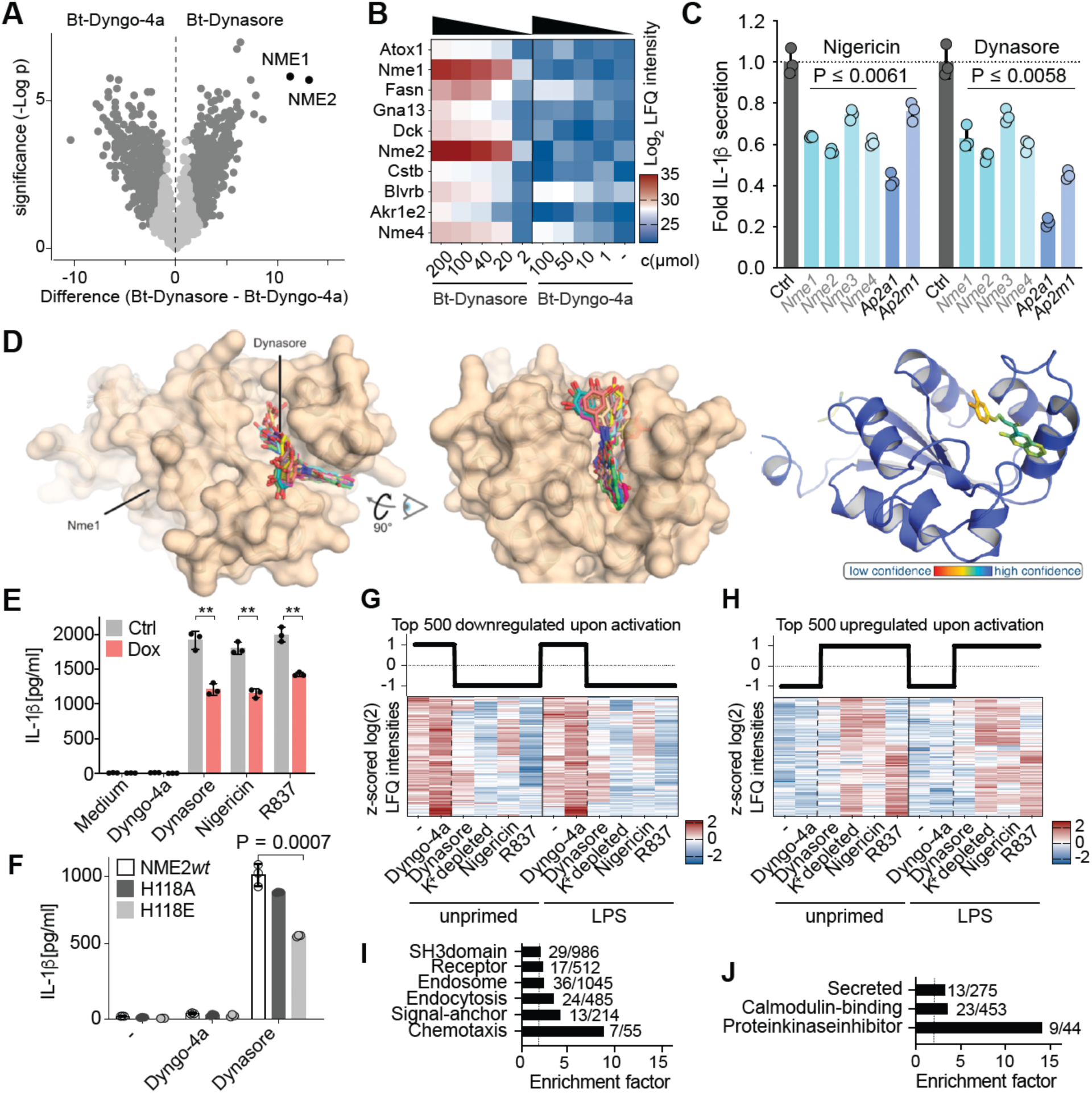
Dynasore interacts with NME1/2 and - like other NLRP3 agonists - activates the NLRP3 inflammasome AP2-dependently. (A) Volcano-plot of streptavidin-bead-based pulldown from LPS-primed ASC^-/-^ iBMDMs treated for 90 min with 100 µM biotin-Dynasore to 100 µM biotin-Dyngo-4a, comparing log2-transformed label-free quantification (LFQ)-intensities student’s t-test fold-enrichment and statistical significance; n = 3. (B) Heatmap showing log2-transformed LFQ-intensities of top-10 enriched interactors in cells treated with biotin-Dynasore compared to biotin-Dyngo-4a across compound concentrations (200, 100, 40, 10, 1, 0 µM biotin-Dynasore; 100, 50, 10, 1, 0 µM biotin-Dyngo-4a). (C) Fold IL-1β release of LPS-primed *wt* Balb/c iBMDMs 48 h after siRNA-mediated knockdown (5 nM) of the indicated genes stimulated with Nigericin (10 µM) or Dynasore (80 µM); n = 3. Statistics indicate significance by student’s two-tailed t-test (P < 0.01)). (D) Structure prediction of Dynasore in complex with hNME1 with Alphafold3. (E) IL-1β release of LPS-primed Balb/c iBMDMs expressing doxycycline-inducible NME1 fused to a mitochondrial targeting sequence peptide (COX8a). Statistics indicate significance by student’s two-tailed t-test (P < 0.01)). (F) IL-1β release of LPS-primed Balb/c iBMDMs expressing dominant negative Histidine118 mutants of NME2 in response to Dyngo-4a (80 µM) or Dynasore (80 µM). Statistics indicate significance by student’s two-tailed t-test; n = 3. (G-J) Phosphoproteomics analysis of inflammasome-stimulated *Nlrp3*^-/-^ iBMDMs. (G) Reference profiles for Pearson’s correlation clustering analysis of log2-transfomred, z-scored 8788 significantly regulated phosphosites across all conditions (One-way ANOVA, S_0_=1, FDR = 0.01) to determine top 500 downregulated phosphosites upon activation as shown in heatmap below. (H) As in (G) for upregulated phosphosites. (I) Enrichment analysis for Uniprot keywords using Fisher exact test on top 500 down- or (J) upregulated phosphosites compared to all 34530 identified phosphopeptides with an enrichment factor > 2.

NMEs are hexameric proteins with pleiotropic functions. They can fuel Dynamin with GTP during the scission of endocytic vesicles (*30*). They can also act as protein kinases and modulate small GTPase activity (*30–32*). As the AP2 complex and its members have been reported to be essential, we depleted AP2 complex subunits as well as NME1-4 by siRNA to elaborate a functional link to NLRP3 activation (Fig. S3C). AP2 is an obligate hetero-tetramer, hence depletion of one subunit results in a destabilization of the whole protein complex. Indeed, we observed reduced inflammasome activation in AP2-and, to a lesser extent, in NME-depleted iBMDMs (Fig. 3C). We further explored whether the local deprivation of NMEs from the plasma membrane inhibits NLRP3 activation. Induced expression of NME1 with a mitochondrial targeting sequence in iBMDMs yielded a reduction of IL-1β release by about 40% across NLRP3 inflammasome agonists (Fig. 3E), pointing to the plasma-membrane localization as a requirement for pathway regulation. Due to the rarity of histidine kinases in eukaryotes (*33*) and its localization within the putative binding pocket of Dynasore, we tested whether NME histidine phosphorylation affects NLRP3 activation. Dominant negative overexpression of a phospho-“dead” H118A and a phospho-“mimetic” H118E of NME2 in iBMDMs alleviated NLRP3 activation significantly (Fig. 3F), but had no impact on BsaK-mediated NLRC4 activation (Fig. S3D). We surmise that both mutants confer important structural changes, suppressing the function of NMEs irrespective of the specific residue. Together our data position AP2 and plasma membrane NMEs in the canonical NLRP3 inflammasome activation pathway.

To establish mechanistic links from a perturbed NME1/2-AP2 axis to known downstream regulations involving GTPase activities, particularly the activation of Rho GTPases and the subsequent phosphorylation of NLRP3 by p21-activated kinases (*34*, *35*), we analyzed NLRP3 induced signaling pathways by phosphoproteomics. Among 34,530 identified and 8,788 significantly changing phosphopeptides across NLRP3 agonists, dephosphorylated pathways (Uniprot keywords) predominated over phosphorylated ones. While proteins involved in kinase inhibition and secretion showed increased phosphorylation (Fig. 3H, J, S3F), we observed inhibited phosphorylation of proteins involved in cell surface signaling, endocytosis, and chemotaxis (Fig. 3G, I, S3E). Specifically, numerous GPCRs, and Rabs including GPR84 and Rab7a, as well as GEF family members such as Dock2, an atypical GEF for Rac1/2 linked to NLRP3 activation were significantly dephosphorylated (Fig. S3E, F). This aligns well with the reported involvement of endocytosis and further points to a potential impairment of cell surface signal transduction upstream of NLRP3. Together, we surmise that NLRP3 activators induce a homeostatic disequilibrium at the cell surface.

### NLRP3 agonists interfere with GPCR signaling and functionally impede directed chemotaxis upstream of inflammasome activation

Inspired by the broad dephosphorylation of cell surface signaling pathways, we next explored how NLRP3 activators functionally affect extracellular signal perception and transduction. We hypothesized that AP2 disruption may affect GPCR signaling due to the prominent role of AP2 in the integration of extracellular cues through receptor mediated endocytosis (*36–39*). Therefore, we quantified whether NLRP3 agonists impact internalization of the β_2_ adrenergic receptor (β_2_-AR) as an exemplary Gs-GPCR via fluorescence microscopy and with a diffusion-enhanced resonance energy transfer (DERET) assay (*40*). We observed rapid β_2_-AR internalization induced by the synthetic agonist isoproterenol, whereas Dynasore, Dyngo-4a or R837 alone did not trigger β_2_-AR internalization (Fig. 4A, B, S4A). Dyngo-4a, Dynasore, and R837 pretreatment diminished isoproterenol-induced β_2_-AR internalization, with Dynasore displaying the strongest inhibition (Fig. 4C, S4B), consistent with its impact on AP2 in addition to Dynamin-inhibition (Fig 4C). To investigate whether GPCRs spatially confined to the plasma membrane by NLRP3 agonists retain their ability to recruit arrestins, we quantified the interaction between β-arrestin2 (β arr2) and isoproterenol-activated β_2_-AR. Indeed, the isoproterenol mediated β arr2: β_2_-AR interaction was preserved for all three agonists, despite differences in recruitment levels (Fig. 4D). We furthermore surveyed isoproterenol-induced βarr2 recruitment to *wt* β_2_-AR in HEK cells, confirming that inflammasome agonists do not prevent β arr2 recruitment (Fig. S4D). Similarly, β-Arrestin 1 (β arr1) recruitment to the CC chemokine receptor type 7 (CCR7), a Gi-coupled GPCR and key regulator of cell migration, was preserved in the presence of NLRP3 agonists after stimulation with its endogenous agonist CCL19 (*41*) (Fig. 4E). As impaired receptor internalization combined with detectable arrestin recruitment is expected to attenuate GPCR signal transduction, we measured cAMP production in HEK293 reporter cells (*42*). Dynasore, Dyngo-4a and R837 strongly reduced norepinephrine (NE)-induced cAMP amplitudes upon β_2_-AR ligation (Fig. 4F), with R837 completely abrogating cAMP production (Fig. S4C). To assess whether signaling inhibition extends to CCR7, we employed a label-free optical biosensor, based on the detection of dynamic mass redistribution (DMR) allowing direct Gi-signaling measurements without Gs-pre-activation (*43*). In accordance with our previous observations, Dynasore, R837, and Dyngo-4a all dampened CCL19-mediated whole cell activation, indicating impaired Gi-GPCR signaling as well (Fig. 4G). Collectively, our data demonstrate that NLRP3 activators disrupt GPCR internalization, attenuate cAMP production, and impair downstream signal transduction consistent with their ability to perturb the AP2 complex.

**Figure 4.**
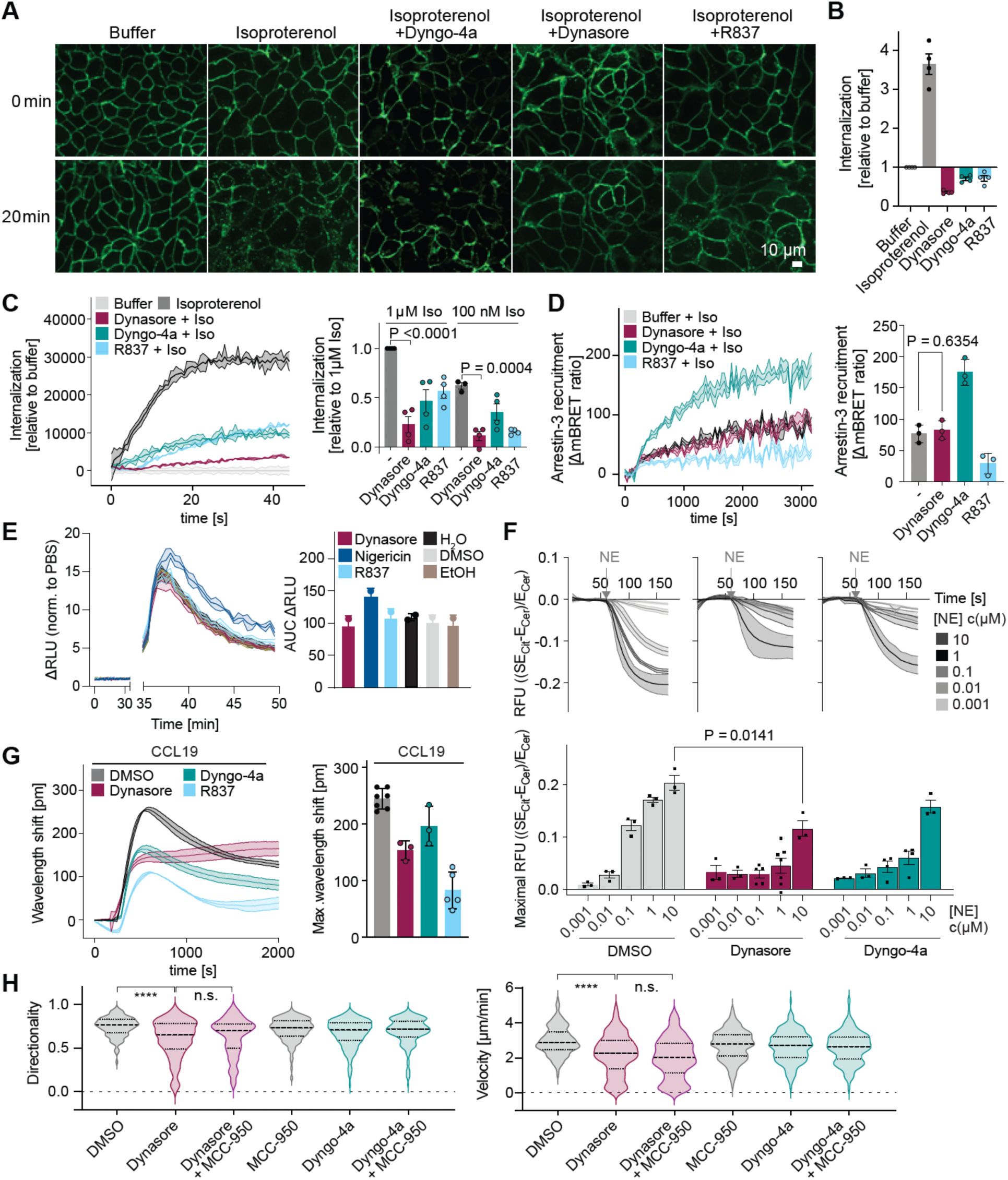
NLRP3 agonists inhibit GPCR internalization, prevent signal transduction downstream of GPCRs and chemotaxis. (A) Fluorescent microscopy of SNAP-β_2_AR-HEK293 cells at baseline (0 min) and 20 minutes after stimulation with 1 µM Isoproterenol +/- 30 µM Dyngo-4a, 60 µM Dynasore or 60 µg/ml R837. (B) Maximum of compound-induced (1 µM Isoproterenol, 60 µM Dynasore, 30 µM Dyngo-4a or 60 µg/ml R837) internalization of overexpressed SNAP-β_2_AR in HEK293 cells relative to buffer between 40 and 50 min measured as diffusion-enhanced resonance energy transfer (DERET). (C) Representative Over-time DERET of isoproterenol (100 nM/1 µM) induced SNAP-β_2_AR internalization in HEK293 cells subsequent to 30 min stimulation with 60 µM Dynasore, 30 µM Dyngo-4a or 60 µg/ml R837(left) and quantification as maximal response relative to buffer between 40 and 50 minutes shown as the mean ± SEM of four independent experiments (right). Statistics indicate significance by student’s two-tailed t-test. (D) Representative Over-time (left) isoproterenol-induced recruitment of β-arrestin 2 to β_2_V_2_ in HEK293 cells measured as BRET between Flag-β_2_V_2_-YFP and Flag-β_2_V_2_-YFP after pre-stimulation with 60 µM Dynasore, 30 µM Dyngo-4a or 60 µg/ml R837or vehicle and quantification as maximal BRET amplitude, shown as the mean ± SEM of three independent experiments (right). Statistics indicate significance by student’s two-tailed t-test. (E) Over-time (left) and maximal (right) CCL19-induced recruitment of β-arrestin 1 to CCR7 in HEK293 cells pre-stimulated 60 µM Dynasore, 10 µM Nigericin, 60 µg/ml R837 or vehicle controls, quantified via bioluminescence resonance energy transfer (BRET). (F) Relative over-time Förster resonance energy transfer (FRET) in HEK293T cells expressing the dual biosensor mlCNBD-FRET after addition of norepinephrine (NE) in the indicated concentrations subsequent to preincubation with either 1% DMSO, 60 µM Dynasore, 30 µM Dyngo-4a; peak cAMP production in response to norepinephrine treatment in indicated concentrations (NE). Cells were pre-treated with DMSO, Dynasore, Dyngo-4a for 30 min before the experiment. (G) Representative Over-time dynamic mass redistribution in HEK293 cells stably expressing CCR7, induced by stimulation with 1 µM CCL19 subsequent to 30 min stimulation with 60 µM Dynasore, 30 µM Dyngo-4a or 60 µg/ml R837 (left) and quantification as the maximum wavelength shift as mean ± SEM of at least three independent experiments. (H) Velocity in [µm/min] and directionality of 50 dendritic cells per replicate during in-vitro 3D collagen migration assay over a timespan of 3 h after treatment with 1% DMSO, 160 µM Dynasore, 10 µM Dyngo-4a or 10 µM MCC-950; n = 4. Statistics indicate significance by Kruskal-Wallis test (n.s. = not significant, **** = P < 0.0001).

GPCRs have widespread functions, thus the physiological consequences of NLRP3-agonist-induced AP2 disruption on GPCR signal transduction are expected to be broad. As NLRP3 activators perturbed chemokine-receptor signal transduction, we evaluated their impact on migratory dendritic cells (DCs). Indeed, Dynasore activated NLRP3 in primary DCs to a similar extent as in BMDMs (Fig. S4E, F). We then examined the functional impact on GPCR-mediated directed chemotaxis. Using a 3D-collagen gel matrix, we analyzed CCL19-induced, CCR7-mediated chemotaxis of DCs and found that Dynasore significantly reduced both the velocity and directionality of these cells, effects that were independent of MCC-950 and therefore upstream of NLRP3 (Fig. 4H). Taken together, our data show that NLRP3 agonists-driven perturbations, particularly AP2 complex disruption, profoundly impair cellular homeostasis by blunting GPCR-mediated integration of extracellular signals, leading to broad physiological consequences.

## Discussion

Our study provides systems-wide insights into organellar and pathway perturbations upstream of NLRP3 inflammasome activation thereby complementing previous spatio-temporal cell biological investigations of NLRP3 activation. We recapitulate organellar disruptions, previously described in the context of NLRP3 activation, such as TGN dispersal, mitochondrial destabilization and a broad impact on endosomes (*9*, *10*, *16*, *44*, *45*). In addition, we identify the dispersal of the AP2 complex as a unifying spatial perturbation shared across NLRP3 agonists. These results are in line with previous findings showing that endocytic disruptions contribute to NLRP3 activation. Our observation that monensin, which acts as a broad organellar disrupting agent but has only mild effects on AP2, can synergistically potentiate K^+^-efflux mediated NLRP3 activation, which depended predominantly on AP2-complex relocalization in K^+^-free HBSS, further refines the recently discussed “two activation signal model” (*10*) for NLRP3 activation. As such, endocytic disruptions may act as recruitment and nucleation sites for NLRP3 and emerge from two central organellar disruption pathways, consisting of the disruption of the mitochondrial electron transport chain and the displacement of AP2. As we did not observe a colocalization between NLRP3 and membrane-depleted AP2 - presumably reflecting “cargo-less pits” (*23*) - full inflammasome activation may occur at sites other than where AP2 loss is initiated. Our experiments identified NME1/2, which supply GTP to the endocytic synapse, as off-targets of Dynasore and revealed a shared AP2 displacement across NLRP3 agonists, while excluding endosomal pH changes or lysosomal leakage. Chemoproteomics and structural predictions suggest that Dynasore directly binds NME1/2 at the plasma membrane. However, because our unbiased discovery approaches inherently detect many regulated proteins, additional contributors cannot be formally excluded. Loss of function studies of the top spatial and chemo-proteomics hits provided causal links between AP2, NME1 and NME2 to NLRP3, however, incomplete inhibition of NLRP3 inflammasome activation suggests functional or spatial redundancy among NMEs. Interestingly, histidine phosphorylation - a very rare signaling moiety in eukaryotes (*46*) at H118 NME1 - contributes to NLRP3 activation, as reported recently for mitochondrial NME4 (*47*). Notably, NME proteins are well conserved, leaving room to speculate that this signaling residue might be involved in danger sensing across organelles or even across the tree of life. Targeting the NME-AP2 axis has many functional implications on cellular homeostasis and our data show that pharmacological interference with NMEs drives cells in a “frozen” state, in line with their formerly proposed role in attenuating cellular motility (*30*). Functional and spatial redundancies of NMEs and lethality of essential gene knockouts like AP2, however, pose conceptual challenges for investigations of this pathway at organism level. The NLRP3 activation-induced broad dephosphorylation of receptor-signaling pathways supports a shutdown of signal transduction. Specifically, Dynasore renders cells unable to receive extracellular cues, as reflected in a general dampening of GPCR mediated cAMP production and reduced directed migration. While endocytosis inhibition per se prevents the uptake of external cues, other NLRP3 agonists such as R837 blocked chemokine stimulated cellular motility even stronger than Dynasore, upstream of pyroptosis. We surmise that this prevention of extracellular signal transduction is due to the spatial deprivation of a key adaptor protein from the plasma membrane and the recruitment of negative regulators of GPCR signaling, as β-arrestin recruitment was preserved to varying degrees by inflammasome agonists. Together, our findings provide mechanistic insight into the spatial and molecular regulation of NLRP3 revealing a fundamental paradigm in which innate immunity senses perturbations of essential cellular pathways to maintain homeostasis.

## Acknowledgments

We would like to thank the Flow Cytometry Core Facility of the Medical Faculty at the University of Bonn for providing support and instrumentation funded by the Deutsche Forschungsgemeinschaft (DFG, German Research Foundation) – Project # 216372545 and Project # 471514137. We further like to thank Victoria Böck, Igor Paron, Mario Oroshi, Nicole Merten, Fabian Paul and Tania Gross for oustanding technical support. Furthermore, we would like to thank the Imaging Core facility of the Institute of Innate Immunity in Bonn, Frank Stein from the Proteomics Core Facility at the EMBL Heidelberg and the Imaging & Protein Production Core Facility of the MPI for Biochemistry.

## Funding

Deutsche Forschungsgemeinschaft (DFG, German Research Foundation) under Germany’s Excellence Strategy – EXC2151 – 390873048 (to F.M.)

German Research Foundation SFB1454 - 432325352 (to F.M.)

German Research Foundation SFB1403 – 414786233 (to F.M.)

German Research Foundation TRR237 – 369799452 (to F.M.)

Swiss National Science foundation (SNSF grant # 220205 to D.F.L.).

## Author contributions

Conceptualization: F.M

Methodology: S.E., K.M.P., Y.A., L.J., F.F., I.S., M.L., M.S.J.M., A.A., N.S., A.F., T.K., J.J.S., B.O.G., D.F., I.B., J.R., J.W., A.K., S.K., L.G., O.J.G., D.I., M.C.T., F.D., R.S., D.W., A.S.

Investigation: S.E., K.M.P., Y.A., L.J., F.F., I.S., M.L., M.S.J.M., A.A., N.S., A.F., T.K., J.J.S., B.O.G., D.F., I.B., J.R., J.W., A.K., S.K., L.G., O.J.G., D.I., M.C.T., F.D., R.S., D.W., A.S.

Visualization: S.E., K.M.P., L.J., F.F., I.S., M.L., M.S.J.M., A.A., N.S., B.O.G., L.G., O.J.G., D.I., G.H., F.M.

Funding acquisition: D.F.L., F.M.

Formal analysis: S.E., K.M.P., L.J., I.S., D.I., G.H.H.B.

Resources; Software: A.U.L., L.N.

Project administration: F.M.

Supervision: M.M.S., G.H.H.B., J.J., G.H., D.F.L., M.M., E.L., E.K., D.W., E.K., F.M.

Writing – original draft: S.E., F.M.

Writing – review & editing: S.E., K.M.P., E.K., G.H.H.B., D.W., E.K., F.M.

## Diversity, equity, ethics, and inclusion

One or more of the authors of this paper self-identifies as a member of the LGBTQ+ community.

## Competing interests

F. M. is a cofounder and consultant of Odyssey Therapeutics. The other authors declare no competing interests.

## Data and materials availability

Our proteomics data is publicly accessible on an interactive website https://immprot.org/nlrp3mapper. The website can be accessed upon request to

We kindly provide any data and scripts for dynamic organellar maps analysis upon request to the authors. Any additional MS data can be accessed upon request to the authors.

Further information and requests for resources and reagents should be directed to and will be fulfilled by the lead contact, Felix Meissner.

## Supplementary Materials

Materials and Methods

Key resource table

Figures S1 to S4

Interactive Website: https://immprot.org/nlrp3mapper

The website can be accessed upon request to

## Materials and Methods

### Experimental model and study participant details

Experiments described in this study were performed with mouse and human primary macrophages, primary mouse dendritic cells, immortalized mouse macrophages and human embryonic kidney cells.

### Animals

Mice were housed under specific-pathogen-free conditions on a 12-hour light/dark cycle in the animal facilities of the Max Planck Institute for Biochemistry or the Institute of Innate Immunity. C57BL/6J wild-type (*wt*) mice were obtained from Jackson Laboratory or Charles River Laboratories. 8–12-week-old male mice were sacrificed by cervical dislocation and used for bone marrow-derived macrophage preparation. Animal experiments were performed according to the German Animal Protection Law.

### Primary cells and cell lines

Immortalized Balb/C *wt*, C57BL/6J *Asc*^-/-^ and *Casp1/11*^-/-^ bone marrow-derived macrophages (iBMDMs), HEK293 as well as HEK293T cells were cultured at 37°C in a humidified incubator under 5% CO_2_ in complete medium (CM), consisting of Dulbecco’s Modified Eagle’s Medium (DMEM; Gibco #10566016) supplemented with 10% heat inactivated calf serum. Cells were cultured up to 80% confluency before passaging and regularly tested for mycoplasma contamination. NME-overexpressing iBMDMs were cultured in CM supplemented with 10 µg/ml blasticidin (InvivoGen, #ant-bl-1). Human monocytes for differentiation into monocyte derived macrophages (MDMs) were obtained from buffy coats from healthy donors according to protocols accepted by the institutional review board at the University of Bonn. Age and Sex of donors was unavailable due to privacy restrictions.

### Method details

#### Generation of mouse bone marrow-derived macrophages

BMDMs were prepared as described elsewhere(*49*). In brief, bone marrow was collected from the femurs and tibiae of C57BL/6J mice and passed through a 70 µm strainer. 5×10^6^ bone marrow cells were plated on non-tissue culture treated, sterile Petri dishes and cultured for 7 days in differentiation media (DMEM GlutaMAX supplemented with 10% heat-inactivated FCS, 5% horse serum, and 20% M-CSF-conditioned medium). Medium was replenished on day 3 of culture. M-CSF-conditioned medium was obtained from L-929 cell supernatants. Cells were detached from plates by incubating in cold phosphate buffered saline (PBS, Gibco #14190094), removed with a cell scraper and replated prior to experiments.

#### Generation of human monocyte-derived macrophages

Buffy coats were diluted 1:1 with PBS and centrifuged over Histopaque-1077 (Sigma, #10771) density gradient medium for 22 min at 900 g. Then, the peripheral blood mononuclear cell (PBMC) fraction was collected and washed three times with PBS, followed by centrifugation for 10 min at 300 g. Monocytes were isolated by negative magnetic selection using human Monocyte Isolation Kit II (Miltenyi Biotec, #130-091-153) following manufacturer’s instructions. 2×10^7^ monocytes were plated on non-tissue culture treated petri dishes for 3 days in RPMI 1640 (Gibco, #61870010) supplemented with 10% FCS, 1% sodium pyruvate (Gibco, #11360070) and 500 U/ml hGM-CSF (ImmunoTools, #11343123). Cells were detached by incubating in cold PBS and replated for experiments.

#### Generation of mouse bone-marrow derived dendritic cells

Bone marrow was collected from 8–12-week-old C57Bl/6J mice. Bone marrow derived dendritic cell (BMDC) differentiation was induced by plating 2×10^6^ cells in 10 ml complete RPMI (RPMI 1640, supplemented with 10% FCS, 100 U/ml penicillin, 100 µg/ml streptomycin (Gibco, #15070063), 50 µM b-mercaptoethanol (Gibco, #21985023)) and 10% GM-CSF containing supernatant from hybridoma culture. Medium was replenished on days 3 and 6 with complete RPMI supplemented with 20% GM-CSF containing hybridoma supernatant. To induce maturation, day 7 or 8 cells were stimulated o/n with 200 ng/ml *E. coli* O111:B4 lipopolysaccharide (LPS) (Sigma-Aldrich, #L4391-1MG).

#### SILAC labelling

iBMDMs were cultured in Dulbecco’s Modified Eagle’s Medium (DMEM) without Arginine, Glutamine, Lysine or Sodium Pyruvate (Gibco, #A14431-01), supplemented with 10% (vol/vol) heat inactivated dialyzed fetal calf serum (PAA, #A11-107), 1 mM Sodium Pyruvate (Sigma, #58636), 1 x GlutaMAX (Gibco, #35050-061) and either; 42 mg/L 13C6,15N4-L-Arginine HCl (Silantes, #201604302) together with 73 mg/L 13C6,15N2-L-Lysine HCl (Silantes, #211604302), or 42 mg/L Arginine HCl and 73 mg/L Lysine HCl with standard isotopic constituents (Sigma, #A6969 and #L8662). Cells were passaged more than 7 times in these media before experiments.

#### Macrophage activation

For spatial proteomics experiments, per replicate, approximately 100×10^6^ *Casp1/11*^-/-^ or *Asc*^-/-^SILAC-heavy or -light labelled iBMDMs were primed with 1 µg/ml LPS-EB or LPS-EB Ultrapure from *E. coli* 0111:B4 (InvivoGen, #tlrl-eblps or #tlrl-3pelps) in CM for 4 h at 37°C, 5% CO_2_, followed by stimulation in serum free conditions with either 10 µM Nigericin (NI; Thermo Fisher Scientific, #N1495), 60 µg/ml R837/Imiquimod (InvivoGen, #tlrl-imqs-1), 10 µM monensin (Merck, #M5273), Mdivi-1 (Selleckchem, #S7162), K^+^-free Hank’s balanced salt solution (HBSS), 160 µM Dynasore (DS; Selleckchem, #S8047), 80 µM Dyngo-4a (DN; Selleckchem, #S7163), or serum-free DMEM as control. Notably, both Dynasore and Dyngo-4a are sequestered by serum proteins and tend to precipitate at concentrations over 200 µM or 100 µM, respectively. We used concentrations which are above the reported maximal IC_50_ for Dynamin-1/2 GTPase inhibition (Dynasore: 15 µM; Dyngo-4a: 2.3 µM).

For phosphoproteomics, per replicate, 1×10^6^ NLRP3-deficient iBMDMs were primed with 1 µg/ml LPS-EB Ultrapure from *E. coli* 0111:B4 in CM for 3.5 h or incubated with CM as control. Then, cells were stimulated with 40 µM Dyngo-4a, 80 µM Dynasore, K^+^-free HBSS, 10 µM Nigericin, 60 µg/ml R837 or serum-free medium as control for 60 min.

For other experiments, 5×10^4^ iBMDMs or 1×10^5^ primary BMDMs seeded one day before the experiment were stimulated with the aforementioned compounds unless indicated otherwise. For NLRC4 activation, Bsak (Florian Schmidt lab) and anthrax protective antigen (PA) (List biosciences, #171E) were mixed in Opti-MEM (Gibco, #31985062) at 2 µg/ml and incubated for 30 min at RT prior to addition to the cells, yielding a final concentration of 1 µg/ml. For experiments to profile AP2a1 relocalization via Western blot, 100×10^6^ *Pycard*^-/-^ iBMDMs were stimulated with 10 µM chloroquine (Sigma, #C6628), 1 µM bafilomycin A1 (Selleckchem, #S1413), 10 µM Wortmannin (Selleckchem, #S2758), 2 mM ATP (InvivoGen, #tlrl-atpl). For experiments in primary BMDMs, BMDCs or iBMDMs which required inhibition of pyroptosis, we used 10 µM MCC-950 (InvivoGen, #inh-mcc), 10 µM VX-765 (InvivoGen, #inh-vx765i-1) or serum-free DMEM as control unless stated otherwise.

#### Enzyme linked immunosorbent assays

Mouse IL-1β, TNF-a and IL-6, as well as human IL-1β in cell supernatants was measured using Enzyme-linked immunosorbent assay (ELISA, DuoSet, #DY401, #DY406, #DY410, #DY201, R&D systems) following the manufacturer’s instructions.

#### Cell death quantification

Cell death was estimated by quantifying the release of lactate dehydrogenase (LDH) into cell supernatants using either CytoTox 96 non-radioactive cytotoxicity assay (#G1780, Promega) or CyQUANT LDH cytotoxicity assay (#C20301, Invitrogen), following the manufacturer’s instructions.

#### Western Blot

For Western Blot analysis of cleaved Caspase-1, 1 ml serum-free cell culture supernatant of inflammasome-stimulated iBMDMs was precipitated by addition of 4 ml 100% ice-cold acetone, followed by three washes with 1 ml 80% acetone and centrifugation at 3000 g for 10 minutes. For analysis of AP2, protein pellets were collected after differential ultracentrifugation. The resulting pellets were dissolved in 1% SDS, 50 mM Tris-HCl pH 7.5. We normalized the input after protein concentration determination using a bicinchoninic acid (BCA) assay (Thermo Fisher Scientific, #23227), followed by reducing and denaturing samples by adding NuPAGE LDS sample buffer (Thermo Fisher Scientific, #NP0007), supplemented with 10 mM DL-Dithiothreitol (Merck, #646563), and boiling for 10 min at 90°C. Proteins were separated by SDS-PAGE on 4-12% Bis-Tris precast gels (Thermo Fisher Scientific, #10247002) using MOPS buffer (Thermo Fisher Scientific, #NP0001) and transferred onto Immobilon-FL PVDF membranes (0.22 µm, Merck, #GVWP04700). Non-specific binding was blocked with 5% non-fat milk in Tris-buffered saline with 0.1% Tween-20 (m-TBST), followed by overnight incubation at 4°C with mouse-anti-mouse Caspase-1 p20 IgG1 (Casper-1, Adipogen, #AG-20B-0042-C100) or mouse-anti-alpha adaptin IgG2a (AC1-M1,1 Abcam, #ab2807), diluted 1:1000 in m-TBST. Membranes were washed three times in TBST for 10 min before incubation in secondary antibody solution (goat anti-mouse IgG H&L, Abcam, #ab97023, 1:5000 for AP2 or donkey anti-mouse IRDYE 800 CW IgG, Li-Cor, #926-32212, 1:20000 for caspase-1, both in m-TBST), three washes in TBST and visualization on a Cytiva or Li-COR Odyssey M system, respectively.

#### ROS measurement

Reactive oxygen species were quantified using CM-H2DCFDA (Thermo Fisher Scientific, #C6827). Briefly, 1×10^6^ *wt* iBMDMs were stimulated with 10 µM Nigericin, 80 µM Dynasore, 40 µM Dyngo-4a, 60 µg/ml R837 or medium for 90 min and washed once with HBSS. 2.5 µM CM-H2DCFDA was added to the iBMDMs and incubated for 30 min 37°C, 5% CO_2_ in the dark. Fluorescence corresponding to ROS production was measured at 494 nm excitation/ 522 nm emission on a FACSCantoä II flow cytometer (BD Biosciences).

### Molecular biology

#### Plasmids and cloning

The Flag-β_2_AR-EYFP plasmid was kindly provided by Prof. Cornelius Krasel (Department of Pharmacology and Toxicology, Philipps University of Marburg, Germany). RFP-tagged-NLRP3 was expressed in a lentiviral vector under a CMV promoter. We cloned eGFP-tagged fusion Ap2a1-Ap2a2 into a lentiviral vector containing a murine phosphoglycerate kinase 1 (PGK1) promoter and a blasticidin selection cassette. The original construct α-eGFP was a gift from Sandra Schmid (Addgene plasmid # 160214; http://n2t.net/addgene:160214; RRID: Addgene_160214). NME1 *wt* was obtained as a string DNA fragment (Thermo Fisher Scientific) and cloned into a lentiviral vector containing a murine PGK1 promoter and a blasticidin selection cassette, or expressed with a C-terminal mitochondrial targeting sequence derived from COX8a under a doxycycline repressor. Single nucleotide exchange on the enzymatic pocket of NME1 (H118>H118A/ H118E) was performed using QuikChange site-directed mutagenesis kit (Agilent, #200519) and verified by sequencing. Cloning of the smBIT-βarrestin1 and βarrestin2 has been described elsewhere (doi: 10.1186/s12964-024-01961-8). The plasmid encoding the β2-adrenergic receptor, which was used in arrestin recruitment experiments was kindly provided by Asuka Inoue (Kyoto University). The plasmid encoding the Nanoluciferase-tagged βarrestin2 (βarrestin2-Nluc) was kindly provided by Carsten Hoffmann (Universitätsklinikum Jena), while the Venus-kRas plasmid was from Nevin Lambert (Augusta University).

#### Lentivirus production and transduction

We used lentiviral transduction in iBMDMs to generate stable cell lines overexpressing NMEs or AP2. For lentivirus production, 2×10^6^ HEK293T cells were transfected with the viral packaging plasmids pMD2.G, psPAX and plasmids harboring the target sequences in a ratio of 1:1.5:2, using GeneJuice (Sigma-Aldrich, #70967) as a transfection reagent. Briefly, lentiviral vectors (3 µg total) and 9 µl GeneJuice were mixed in 300 µl Opti-MEM, incubated for 20 min at 21°C and added on to HEK293T cells, followed by incubation for 6 h at 37°C, 5% CO_2_. Transfection medium was exchanged with CM and cells were incubated for 3 days at 37°C, 5% CO_2_ for virus production. Viral supernatants were harvested by aspiration with a blunt needle, cleared from debris by passing through a 0.45 µm PTFE syringe filter (Merck Millipore, #SLFH050) and added to 1×10^6^ *wt* iBMDMs for overnight transduction. Blasticidin antibiotic selection was started 2 days after transduction and iBMDMs were passaged at least twice before proceeding with experiments. Generation of an isogenic HEK293 cell clone transfected to stably express an N-terminally SNAP-tagged β_2_-adrenergic receptor (SNAP-β_2_AR) was carried out in collaboration with Prof. Hanns Häberlein (Department of Biochemistry, Bonn University, Germany), as previously reported(*50*).

#### siRNA-mediated gene knockdown

Gene knockdown by small interfering RNA (siRNA) transfection was performed using a pool of 4 siRNAs per gene (Dharmacon, Ap2a1 #L-043307-01-0010, Ap2m1 #L-042924-01-0010, Nme1 #L-040142-00-0005, Nme2 #L-040143-00-0005, Nme3 #L-049492-00-0005, Nme4 #L-049846-00-0005, Pycard #L-051439-01-0010, Casp1 #L-048913-00-0005, Casp11 #L-042432-00-0005, Aim2 #L-044968-00-0005, Nlrp1a #L-066229-00-0005, Nlrc4 #L-055000-00-0005, non-targeting control #D-001810-10-05). In brief, 50 nM siRNA were mixed with 15 µl lipofectamine RNAiMAX (Thermo Fisher Scientific, #13778075) in 600 µl Opti-MEM (Gibco, #31985062), incubated for 5 min at 21°C, before addition of 2×10^6^ *wt* iBMDM in 2 ml serum free DMEM on top. Cells were incubated for 6 h, 37°C, 5% CO_2_ and supplemented with 3 ml CM. After 48 h incubation, cells were plated for experiments.

#### Potassium quantification by atomic absorption spectrometry

We determined intracellular K^+^ in macrophages from 3 C57Bl6 *wt* BMDMs as previously described(*51*). Briefly, 1 million BMDMs were primed with 1 µg/ml LPS in CM and stimulated with 80 µM Dynasore, 10 µM Nigericin ± 10 µM MCC-950 in serum-free DMEM supplemented with GlutaMax, Pyruvate and Non-essential amino acids or incubated in medium alone. Cells were then washed in osmolarity-matched sucrose solution (290 mOsm/kg) for a total of three times and lysed in 100 µl 0.1% Triton-X in ddH_2_O on ice for 30 min. K^+^ content was determined using an iCE 3000 Series atomic absorption spectrometer (Thermo Fisher Scientific).

#### Internalization analysis via diffusion-enhanced resonance energy transfer (DERET)

The ability of Dynasore (60 µM), Dyngo-4a (20 µM), and R837 (60 µg/ml) to induce SNAP-β_2_AR internalization was measured as previously described^40,^(*52*, *53*). Briefly, stable SNAP-β_2_AR HEK293 cells(*50*) were seeded into pDL-coated, white, flat bottom 96-well plates (Sarstedt, order-No. 82.1581.210) at 50000 cells per well and cultivated for 24 hours at 37 °C and 5 % CO_2_. Cells were labelled with 50 µl of 100 nM SNAP-Lumi4-Tb (Revvity, #SSNPTBC) in HBSS supplemented with 20 mM HEPES for one hour at 37 °C. After removal of labeling reagent, cells were washed four times with assay buffer. Then, 50 µl fluorescein (Sigma-Aldrich, #2321-07-5) diluted in assay buffer was added to reach a final concentration of 25 µM. Thereafter, Dynasore, Dyngo-4a, and R837 were added and incubated for 30 minutes at 37 °C, 5 % CO_2_. Donor (Lumi4-Tb, 620 nm) and acceptor (fluorescein, 520 nm) emissions were measured after excitation at 337 nm using a PHERAstar FSX multimode plate reader (BMG LabTech) pre-heated to 37 °C. After a baseline read, isoproterenol diluted in a volume of 60 µl was added at various concentrations. Internalization was measured for 50 minutes and quantified by calculating the donor/acceptor ratio using MARS data analysis software (BMG Labtech, Version).

#### Arrestin recruitment analysis via bioluminescence resonance energy transfer (BRET)

Recruitment analysis of β-arrestin 2 to β_2_V_2_, (a β_2_ receptor variant harboring the vasopressin V2 receptor C-terminus to enhance arrestin interaction sensitivity) was performed as previously described(*54*). Briefly, 2×10⁶ HEK293 cells were transiently transfected 48 h before the experiment with 500 ng RLuc-arr3 and 500 ng Flag-β_2_V_2_-YFP and 4000 ng of DNA with pcDNA3.1(+) using polyethylenimine (PEI, 1 mg/ml) at a DNA to PEI ratio of 1:3. The following day, cells were detached and seeded at a density of 100,000 cells per well in white 96-well plates. Then, coelenterazine was added at a concentration of 5 µM and incubated for 15 minutes at 37 °C. RLuc-luminescence (475 nm ± 30 nm) and YFP-fluorescence (535 nm ± 30 nm) were measured for 1.0 s every 40 s for 10 minutes at 37 °C using the PHERAstar® FSX plate reader (BMG Labtech). The BRET ratio was calculated using MARS data analysis software (BMG Labtech, Version 3.3.2) by dividing the acceptor emission by the donor emission and multiplying by 1000.

Furthermore, recruitment of βarrestin2 to the wild-type β_2_-adrenergic receptor (β_2_AR) was measured using a bystander BRET assay setup. Here, the BRET acceptor (Venus) is confined to the plasma membrane (Venus-kRas), while the BRET donor (Nanoluciferase or Nluc) is fused to the C-terminus of βarrestin2 (βarrestin2-Nluc). Upon receptor activation, βarrestin2-Nluc, which is cytosolic in the basal state, translocates to the plasma membrane leading to an increase in BRET to Venus-kRas. HEK293T cells (2×10^6^ cells in 9 mL) were transfected in suspension with 900 ng of β_2_AR, 90 ng of βarrestin2-Nluc and 2250 ng of Venus-kRas. Empty pcDNA3.1 was used to adjust the amount of transfected DNA to 9000 ng. Transfected cells were then seeded (100 µL/well) into a poly-D-lysine-coated, white, opaque 96-well plate and incubated in a water-saturated atmosphere (37 °C, 5% CO_2_). After two days, cells were washed once with assay buffer (Hank’s Balanced Salt Solution (HBSS) supplemented with 20 mM HEPES) and incubated with Dyngo-4a (20 µM), Dynasore (60 µM), R837 (60 µg/mL) or DMSO (all dilutions prepared in assay buffer) for 15 min at 37 °C. Next, Nano-Glo substrate (Promega, N1120, final dilution: 1/1000) was added to the cells. After an additional incubation for 15 min at 37 °C, three baseline reads were recorded, followed by the addition of a serial dilution of isoproterenol. BRET measurements were performed at 37 °C using the BRET1plus module (donor: 475 ± 30 nm; acceptor: 535 ± 30 nm) installed in a Pherastar FSX plate reader (BMG Labtech). BRET was defined as the ratio of sensitized acceptor emission over donor bioluminescence. For all wells and timepoints, the obtained (raw) BRET ratios were corrected for interwell variability by subtracting the average of the baseline values (separately for each well) followed by a subtraction of the average of the buffer values. Maximal values of the traces were used to construct concentration-response curves for each independent experiment.

#### Dynamic mass redistribution (DMR) analysis

Label-free whole cell biosensing based on detection of dynamic mass redistribution (DMR) was performed as previously described in detail^43,^(*55*). Briefly, 18000 HEK293 cells per well were seeded onto fibronectin-coated 384-well Epic biosensor plates (Corning, #4585) and cultured at 37 °C in a humidified atmosphere with 5% CO₂. After 24 hours, the medium was replaced with Hanks’ Balanced Salt Solution (HBSS, with or without potassium) supplemented with 20 mM HEPES and 0.45% DMSO to ensure matched DMSO concentrations between the medium and the added compounds. Cells were then equilibrated in an Epic Reader (Corning) at 37 °C for 60 minutes. Following equilibration, Dynasore (60 µM), Dyngo-4a (20 µM), and R837 (60 µg/ml) were added for a duration of 30 minutes. Data acquisition was performed for an additional 60 minutes post-compound addition using the semi-automated Cybio-Selma liquid handling system (Analytik Jena). Results are expressed as buffer-corrected mean picometer (pm) shifts ± SD over time. Quantification was performed by calculating either the maximum pm shift or the area under the curve (AUC) over time for each experiment, reported as mean ± SEM from three independent biological replicates.

#### Real-time, FRET based single cell cAMP detection

Live-cell cAMP measurements were performed using HEK293 cells expressing the mICNBD-FRET cAMP sensor. Cells were seeded at a density of 3×10^4^ cells per well of a black PhenoPlate 96-well plate (Revvity, #6055302) and incubated at 37°C and 5% CO_2_ for 24 h before the measurement. For cAMP measurements, the cells were washed once with 200 µl extracellular solution (ES, 10 mM HEPES pH 7.4, 120 mM NaCl, 5 mM KCl, 2 mM MgCl_2_, 2 mM CaCl_2_, 10 mM glucose), followed by addition of 100 µl ES into each well. Then, 100 µl of 2x stock solutions (120 µM Dynasore, 60 µM Dyngo-4a, 60 µM Barbadin, 60 µg/ml R837 or 2% DMSO) were either added acutely for direct cAMP measurements or 30 min before as pre-incubation. Imaging was performed using the Zeiss Axio Observer Z1 with a dual camera setup operating at 37°C, using the ZEN software (Ver. 2.6.76.00000). FRET was recorded by exciting cerulean at 436 nm and measuring the emission of cerulean and citrine at 470 nm and 535 nm, respectively. The measurements were performed with 10 second intervals, with the first 6 images (60 s) serving as the baseline, followed by the addition of the stimulants at the indicated timepoints. To measure the cAMP increase in response to the indicated agonists, cells were stimulated with varying concentrations of norepinephrine (NE, Sigma, #E4642), followed by addition of 40 µM Forskolin (Sigma, #f6886) as a positive control. The change in FRET was calculated using the sensitized emission (SE) of citrine and cerulean (. Data were collected using Microsoft Excel (Ver. 16.88) and analyzed with GraphPad Prism (Ver. 10.3.0). The first 6 images were used as the baseline. The fractional difference (of inverse values, normalized to 𝐸 the 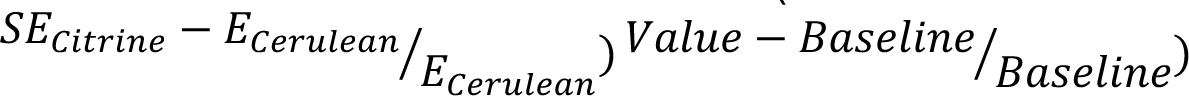 smallest and highest value within the data set was calculated for plotting. For the dose response curves, the maximal amplitudes of the NE response for each condition were determined and min-max normalized from highest to lowest concentration.

#### In vitro 3D collagen migration assay

For 3D in vitro migration, 2×10^5^ mature BMDCs were resuspended in complete RPMI and mixed with a collagen suspension to a final collagen concentration of 1.66 mg/ml (Cellsystems, #5005-100ML), supplemented with 1x minimum essential medium eagle (MEM, Gibco, #11095080) and 0.4% sodium bicarbonate (Sigma, #S6014). BMDC-collagen mixtures were cast into custom-made migration chambers as described elsewhere(*56*, *57*) and incubated for 1 h at 37°C, 5% CO_2_ to engage polymerization. CCL19 (R&D Systems, #440-M3) was suspended to 100 nM in complete RPMI and placed on top of the gel. Migration chambers were sealed with paraffin wax (Sigma, #411663) and unpolymerized gels were excluded from the study. Stimuli (160 µM Dynasore, 10 µM Dyngo-4a or 30 µg/ml R837) were added to BMDCs during gel polymerization and during image acquisition together with CCL19. Where applicable, BMDCs were pre-treated with 10 µM MCC-950 for 30 min. Image acquisition was performed with a Nikon widefield microscope and a C-Apochromat 10x/0.3 PH1 air objective and the software NIS-elements 6.0 (Nikon). Images were acquired in 120 s intervals for 3 h at 37°C, 5% CO_2_. 50 cells were tracked manually using the ‘Manual tracking Plug-in’ for ImageJ (ImageJ2, Ver. 2.14.0/1.54f). The ImageJ Chemotaxis tool was used to determine average (frame-to-frame) speed and directionality as per Euclidean distance from start to end of the track/ total track distance.

#### β_2_AR-Internalization via microscopy

Fluorescence microscopy of stable SNAP-β_2_AR HEK293 cells was performed on a Zeiss Axio Observer Z (Carl Zeiss, Jena, Germany), equipped with ApoTome 2.0 using 63x magnification. Definite Focus (Carl Zeiss, Jena, Germany) was applied to maintain focal plane stability. Cells were seeded into a pDL-coated eight-well microscopy plate (Ibidi, order-no. 80826) at a density of 120000 per well and cultivated for 24 hours at 37 °C and 5 % CO_2_. Following three washes with HBSS, 200 μl of HBSS supplemented with 20 mM HEPES and 50 µl solution containing Dynasore (60 µM), Dyngo-4a (20 µM) or R837 (60 µg/ml) was added to each well. Baseline images were recorded immediately before stimulation with 50 μl Isoproterenol. Subsequent images were collected at specified time points. Image analysis was conducted using Zen blue Imaging software (Carl Zeiss, Jena, Germany).

#### Tagged NLRP3 and AP2A1 imaging

For imaging of AP2a1-NLRP3 to analyze colocalization, *wt* iBMDMs transiently transfected with plasmids encoding AP2a1-AP2a1-eGFP fusion and RFP-tagged-NLRP3. For reverse transfection, per well, 0.5 µl lipofectamine 2000 (Thermo Fisher Scientific, #11668027) were mixed with 50 ng plasmid DNA (25 ng each) in 50 µl Opti-MEM and incubated for 20 min at 21°C. 5×10^4^ iBMDMs per well were added on top in a black PhenoPlate 96-well plate (Revvity, #6055302) and incubated at 37°C and 5% CO_2_ for 24 h before the measurement. Cells were primed with 1 µg/ml LPS for 3.5 h and stimulated with 80 µM Dynasore, 40 µM Dyngo-4a, 10 µM Nigericin or serum-free medium in presence of 10 µM VX-765 for 90 min. After washing with 1x PBS, cells were fixed in 4% paraformaldehyde (PFA, Thermo Fisher Scientific, #28906) in PBS for 20 min at 37°C, washed twice with PBS and kept in 100 µl PBS until imaging. Wells were imaged using a Leica SP8 lightning confocal microscope at 63x magnification.

#### ASC specking analysis in BMDCs

BMDCs were transferred onto coverslips, incubated 5- 10 min at 37°C to adhere and fixed with 4% formaldehyde (Thermo Fisher Scientific, #28908) for 20 min. Fixed cells were washed twice with PBS. For immunofluorescence staining, cells were permeabilized with 0.2% Triton-×100 (Sigma, #T8787) in PBS for 30 min and blocked with 1%BSA (Roth, #1ETA.3) in PBS for 1 h at RT. Cells were incubated with anti-ASC (Merck, #04-147, 1:1000 in blocking solution) o/n at 4°C. Cells were washed three times with PBS and incubated for 1 h with donkey anti-mouse Alexa Fluor 647 Affinipure F(ab’)2 fragment IgG (H+L)(Jackson ImmunoResearch, #715-606-150, 1:400 in PBS), followed by three more washes. Confocal microscopy was performed using a Zeiss LMS880 Confocal Laser Scanning Microscope with a 40x objective, using the imaging software ZEN Black 2.3 SP1. Downstream image analysis was performed with ImageJ.

### Sample preparation for LC-MS/MS

#### Full proteomes

Total lysates were diluted to a final urea concentration of 2 M and sonicated on ice for 15 min (level 5, Bioruptor, Diagenode). Cell supernatants (400 μl each) were denatured with 2 M urea in 10 mM HEPES (Gibco, #15630080), pH 8. Proteins of both sample types were reduced with 10 mM dithiotreitol (DTT, Sigma, #646563) for 30 min at RT followed by alkylation with 55 mM iodoacetamide (IAA, Sigma, #I6125) for 20 min at RT in the dark. Remaining IAA was quenched with 100 mM thiourea. Proteins were digested with 1 μg LysC (#129-02541, Wako Chemicals) at RT for 3 h and 1 μg trypsin (#T6567, Sigma) at RT overnight. Protein digestion was stopped with 0.6 % trifluoroacetic acid and 2 % acetonitrile before peptides were loaded onto reversed phase C18 StageTips (#2215, 3M^TM^ Empore^TM^, IVA Analysentechnik). Supernatants and 50 μg of total lysates were loaded onto the C18 StageTips. Peptides were desalted using 0.5 % acetic acid and subsequently eluted from the C18 StageTips with 50 μl 80 % acetonitrile in 0.5 % acetic acid. After concentrating and drying in a SpeedVac (Thermo Fisher Scientific), peptides were resuspended in 10 μl 2 % acetonitrile, 0.1 % trifluoroacetic acid and stored at -20°C until mass spectrometric analysis.

#### Streptavidin-bead affinity purification

Per replicate, 5×10^6^ ASC-deficient iBMDMs were primed with 1 µg/ml LPS-EB Ultrapure from *E. coli* 0111:B4 in CM for 3.5 h, rinsed twice with PBS, harvested by scraping, pelleted by centrifugation at 300 g for 5 min and snap frozen in liquid nitrogen. The resulting pellets were thawed on ice and resuspended in 500 µl lysis buffer, consisting of 1 mM CaCl_2_, 0.5 mM MgCl_2_, 140 mM NaCl, 1% NP-40 (Thermo Fisher Scientific, #85125), supplemented with 7.5 U/ml smDNAse (MPI for Biochemistry) and cOmplete EDTA-free protease inhibitor (Roche, # 11836170001), followed by incubation for 1 h at 4°C while shaking on a Thermomixer (Eppendorf) at 500 rpm. The lysates were then incubated with either biotin-Dynasore (891.05 g/mol, 200 µM, 100 µM, 40 µM, 10 µM, 1 µM), biotin-Dyngo-4a (907.05 g/mol, 100 µM, 50 µM, 10 µM, 1 µM) or DMSO as vehicle control for 2 h at 21°C while shaking at 500 rpm. The lysates were then cleared by centrifugation at 10000 g for 10 min at 4°C and 300 µl of cleared supernatant were used for affinity purification. Next, 50 µl streptavidin-agarose (Thermo Fisher Scientific, #20359) per sample were equilibrated in equal volumes lysis buffer, pelleted by centrifugation at 100 g for 5 min and added to each sample, followed by incubation for 2 h at 4°C while shaking at 2000 rpm. Then, the beads were pelleted by centrifugation at 300 g for 3 min, 4°C and the supernatant was discarded. The beads were washed twice in lysis buffer, then washed twice in 40 mM HEPES, followed by on-bead lysis with 50 µl 6 M Urea in 40 mM HEPES on ice for 20 min before reduction and alkylation with 10 mM DTT, 50 mM IAA and digesting over night with 1 µg Trypsin/ Lys-C mix (Promega, #V5072) in 2 M Urea, 50 mM Tris pH 8.5 at 21°C while shaking at 800 rpm. Peptides were cleaned up using C18 stage tips.

#### Spatial proteomes

Dynamic organellar maps were generated as described elsewhere(*14*). Briefly, stimulated macrophages were washed twice in cold isotonic lysis buffer (ILB, 120 mM glycerol, 140 mM KCl, 25 mM Tris pH 7.5, o.2 mM EGTA, 0.5 mM MgCl_2_ supplemented with protease inhibitors), harvested using a cell scraper and centrifuged at 400 g, 5 min before aspiration of the supernatant and kept on ice until lysis. Cells resuspended in 3.5 ml ILB were passed 10-13 times through a preequilibrated cell homogenizer (Isobiotec) with a 10 µm clearance ball to generate a crude lysate. Intact cells were then depleted by centrifugation at 300 g, 5 min. The supernatant was then subjected to differential ultracentrifugation using an Eppendorf centrifuge and a Beckman Coulter Optima-Max ultracentrifuge at 4°C, starting at 1000 g for 10 min, 3000 g for 10 min, 5400 g for 15 min, 12200 g for 20 min, 24000 g for 20 min and lastly 78400 g for 30 min, all while transferring the supernatant to a fresh (ultracentrifuge-)tube and resuspending the resulting pellets in membrane re-solubilization buffer (1% sodium deoxycholate (SDC, Sigma, #D6750), 1% Igepal CA-630 (Sigma, #I8836), 0.1% sodium dodecyl sulfate (SDS, Sigma, #436143), 150 mM NaCl, 25 mM Tris pH 8, supplemented with protease inhibitors). Protein concentration was determined using BCA. 30 µg of each SILAC light-labelled fraction were independently mixed with an equal amount of SILAC heavy reference organellar fraction. Protein was precipitated by addition of 5 volumes of ice-cold acetone, before overnight incubation at −20°C and subsequent centrifugation at 4°C for 5 min at 10000 g. In case of nuclear, organellar, and cytosol fractions, 60 µg of SILAC light labelled sample only were subjected to the same precipitation regime. All subsequent steps were performed at room temperature. Supernatants were removed and pellets allowed to air-dry for 5 min. Pellets were re-suspended in digestion buffer (50 mM Tris pH 8.1, 8 M Urea, 1 mM DTT) and incubated for 15 min. Cysteines were alkylated by addition of 5 mM Iodoacetamide with incubation for 20 min. Proteins were enzymatically digested by addition of 1 µg LysC per 50 µg of protein, with incubation for 3 h. Digests were then diluted four-fold with 50 mM Tris pH 8.1 before addition of 1 µg Trypsin per 50 µg of protein and overnight incubation.

Peptides were fractionated using stage tips with 2X discs of SDB-RPS material (3M^TM^ Empore^TM^, IVA Analysentechnik, #2241). Stage tips were activated with 100% Acetonitrile, followed by an aqueous solvent containing 30% Methanol and 1% TFA. Peptide mixtures were acidified with 1% TFA, 15 µg (for single shot) or 25 µg (for fractionation) was loaded onto activated stage-tips, and washed with 0.1% TFA. Peptides were eluted with 60 µl buffer X (80% Acetonitrile, 5% Ammonium Hydroxide) for single shot analyses. For fractionation, peptides were eluted using 20 µl SDB-RPSx1 (100 mM Ammonium formate, 40% Acetonitrile, 0.5% Formic acid), then 20 µl SDB-RPSx2 (150 mM Ammonium formate, 60% Acetonitrile 0.5% Formic acid), then 30 µl buffer X. Tryptic peptides were dried almost to completion in a centrifugal vacuum concentrator (Concentrator 5301, Eppendorf), and volumes were adjusted to 10 µl with buffer A* (0.1% TFA, 2% Acetonitrile).

#### Phosphoproteomes

Cells were washed twice in regular or K^+^-free HBSS and lysed on ice for 20 min with 100 µl 8 M Urea in 50 mM Tris pH 8.5, containing smDNAse and PhosSTOP (Roche, # 4906845001). Lysates were cleared by centrifugation at 10000 g for 10 min at 4°C. Protein concentration was equalized via BCA, Urea was diluted to < 2 M with 50 mM Tris pH 8.5 and proteins were reduced and alkylated using 10 µM Tris(2-carboxyethyl)phosphine (TCEP, Thermo Fisher Scientific, #77720) and 50 µM 2-chlroacetamide (CAA, Sigma, #C0267) for 30 min at 21°C, 800 rpm using an Eppendorf Thermomixer. Proteins were cleaned up prior digestion and phosphopeptides enrichment via protein aggregation capture (PAC) as described elsewhere(*58*). Briefly, per 20 µg protein, 5 µl MagReSyn hydroxyl beads (Resyn biosciences, #MR-HYX005) were equilibrated and washed twice with 70% acetonitrile. Protein extracted were added to the beads and adjusted to 70% acetonitrile, followed by 10 min on-bead precipitation. Then, the beads were washed with 100% acetonitrile, 70% ethanol and finally resuspended in digestion buffer (50 mM Tris pH 8.5, 1 µg Trypsin/ Lys-C mix per 50 µg protein) to digest overnight at 37°C. Digestion was stopped by acidification with 1.5% formic acid. Phosphopeptides were enriched using the AssayMAP Bravo liquid handling platform with 5 µl Fe(III)-NTA cartridges (AssayMAP, #G5496-60085), following the manufacturer’s instructions. Lastly, phosphopeptides were desalted for LC-MS/MS using Evotips (Evosep, #EV2011).

#### LC-MS/MS

Peptide mixtures were analyzed in a single-run liquid chromatography mass spectrometry (LC-MS/MS) format. Each peptide mixture was loaded onto a C18-reversed phase column (50 cm, 75 μm inner diameter) and separated with a non-linear gradient of 2 - 60 % buffer B (80 % acetonitrile, 0.1 % formic acid) at a flow rate of 250 nl/min over 120 min for global- and phospho-proteomes and 180 min for SILAC maps using a nanoflow UHPLC instrument (Easy-nLC 1200, Thermo Fisher Scientific). Chromatography columns (#TSP075375, Composite Metal Service Ltd.) were in-house packed with ReproSil-Pur 120 C18-AQ 1.9 μm resin (#r119.aq., Dr. Maisch GmbH) in methanol. Chromatography and column oven (Sonation GmbH) temperature were controlled and monitored in real-time with SprayQC. Column oven temperature was set to 50°C. Separated peptides were analyzed on benchtop quadrupole-Orbitrap instruments (Q Exactive HF/HFx mass spectrometer, Thermo Fisher Scientific or Orbitrap Exploris 480 mass spectrometer, Thermo Fisher Scientific) or with a nanoelectrospray ion source (Thermo Fisher Scientific), which was coupled on-line to the liquid chromatography instrument. For full and SILAC spatial proteomes, the mass spectrometer was operated in a data dependent mode with a survey scan range of 300-1650 m/z and a resolution of 60,000 at m/z 200. Up to the 10 most abundant precursor ions with charge states 2 to 5 were isolated for higher-energy collisional dissociation (HCD) with isolation windows of 1.8 m/z. Normalized collision energies (NCE) for HCD were set to 26. Fragmentation spectra were acquired with a resolution of 15,000 at m/z 200. Dynamic exclusion duration of sequenced peptides was set to 30 s to reduce repeated peptide sequencing. Maximum ion injection times were 20 ms for the full MS scan and 55 ms or the MS/MS scan. Automatic gain control (AGC, ion target values) was set to 3e6 for the survey and 1e5 for the MS/MS scan. For phosphoproteomics, peptide mixtures were analyzed in a single-run liquid LC-MS/MS format. Each peptide mixture was loaded onto a 3^rd^ generation AuroraElite HPLC column (15 cm, 75 µm inner diameter, filled with C18 resin with a bead size of 1.7 µm, IonOpticks, #AUR3-15075C18), connected to an Evosep One (Evosep) with a non-linear gradient of incremental buffer b (99.9% acetonitrile, 0.1% formic acid) using the whisper zoom 40 SPD method at a flow rate of 200 nl/min over 32.5 min. Chromatography and column oven (Sonation GmbH) temperature were controlled and monitored in real-time with SprayQC. Column oven temperature was set to 50°C. Separated peptides were analyzed on a benchtop quadrupole-Orbitrap Astral mass spectrometer (Thermo Fisher Scientific). The mass spectrometer was operated in data independent acquisition mode with a survey scan range of 380-980 m/z and a resolution of 240000 at m/z 200, with a maximum inject time of 5 ms. Peptides within a 2 m/z window size were isolated for HCD with NCE set to 27%. MS/MS scan range was set to 150-2000 m/z with a maximum inject time of 3.5 ms. Automatic gain control was set to 300% for survey and 5e4 or 500% for the MS/MS scan.

#### Quantification and statistical analysis

Analysis of DDA MS raw files for spatial proteome data, affinity purification MS and full proteomes was performed using the proteomics platform MaxQuant (Versions 1.6.2.1, 2.0.3.0). Files were searched for tryptic peptides with a minimum length of 7 amino acids against the mouse SwissProt canonical and isoform protein database (UniProt) and a contaminant database within the Andromeda search engine. Cysteine carbamidomethylation was set as fixed and methionine oxidation and N-terminal acetylation as variable peptide modifications. False discovery rates (FDRs) were set to 1% for peptide and protein level identification with allowed initial protein mass deviation up to 4.5 ppm and maximum fragment mass deviation of 20 ppm. We used the “match between runs” feature (match time window size = 0.7 min, alignment time window = 20 min) for nonlinear retention time alignment in MaxQuant and performed label-free quantification (LFQ) via MaxLFQ at a minimum ratio count of 1. For fractionated spatial proteomes, identical centrifugation fractions were collapsed to one sample by setting the same experiment name. For SILAC data, multiplicity was set to 2, with Lys8 and Arg10 as heavy labels and enabled Re-quantify feature with a minimum number of quantification events of 2.

Data filtering, normalization and analysis was performed using the R and R studio environment for statistical data processing or the Perseus platform (Version 1.6.15, 2.0.3). Data were visualized with the package “ggplot2” or using GraphPad Prism (Version 10). Data filtering and normalization were performed as described previously using customizable R scripts.

For detection of proteins undergoing significant changes in localization, a simple subtraction between the normalized profiles of treated vs untreated samples per fraction/ speed was calculated to generate a delta matrix of delta profiles for each protein as described previously. The Mahalanobis distance to the center of the distribution of delta values was calculated for each protein, using the minimum covariance determinant method, implemented in the Perseus software. After conversion of distances to p-values, Benjamini Hochberg correction for multiple hypothesis testing was performed. Proteins with corrected p-values below 0.05 in at least 2 out of 3 experiments, and high correlations above 0.7 in delta profiles between replicates show a significant change in localization upon treatment. The absolute value of the delta values for each fraction were summed to generate a metric referred to as “absolute distance”. In order to detect significantly changing protein categories, proteins were annotated with GO terms and a Fisher exact test was performed on outliers or on an absolute distance cut-off correlating with outliers. Phosphoproteomics DIA data were analyzed using Spectronaut (Version 19.7.250203). Class I phosphosites with a localization probability > 0.75 were filtered for 100% valid values in at least one group (according to treatment), log_2_-transformed and subjected to column-wise z-score normalization. Missing values were imputed from a normal distribution with a downshift of 1.8, width of 0.3. To determine significantly changing phosphosite intensities, an ANOVA across all conditions was performed with a S_0_ fudge factor of 1 at a FDR or 0.01. Significantly changing phosphosites were further surveyed for shifts according to treatment (up- or downregulated during activation, Fig. 3G, H) and the resulting profiles were analyzed for changing categories using Keywords annotations in the built-in Fisher Exact test.

To determine global changes between biotin-Dynasore/ biotin-Dyngo-4a-treated vs untreated controls, LFQ intensities were grouped (triplicate control vs triplicate treatment experiments), log 2 transformed, and filtered for a minimum of three valid values in at least one group. Missing values were then imputed from a normal distribution (downshift = 2, width = 0.3). A two-tailed Student’s t-test was performed between control and treatment for each protein with a permutation-based correction for multiple hypothesis testing. Significantly changing protein categories were detected using the 1D annotation enrichment analysis for UniProt keywords terms on the fold changes between control and treated conditions.

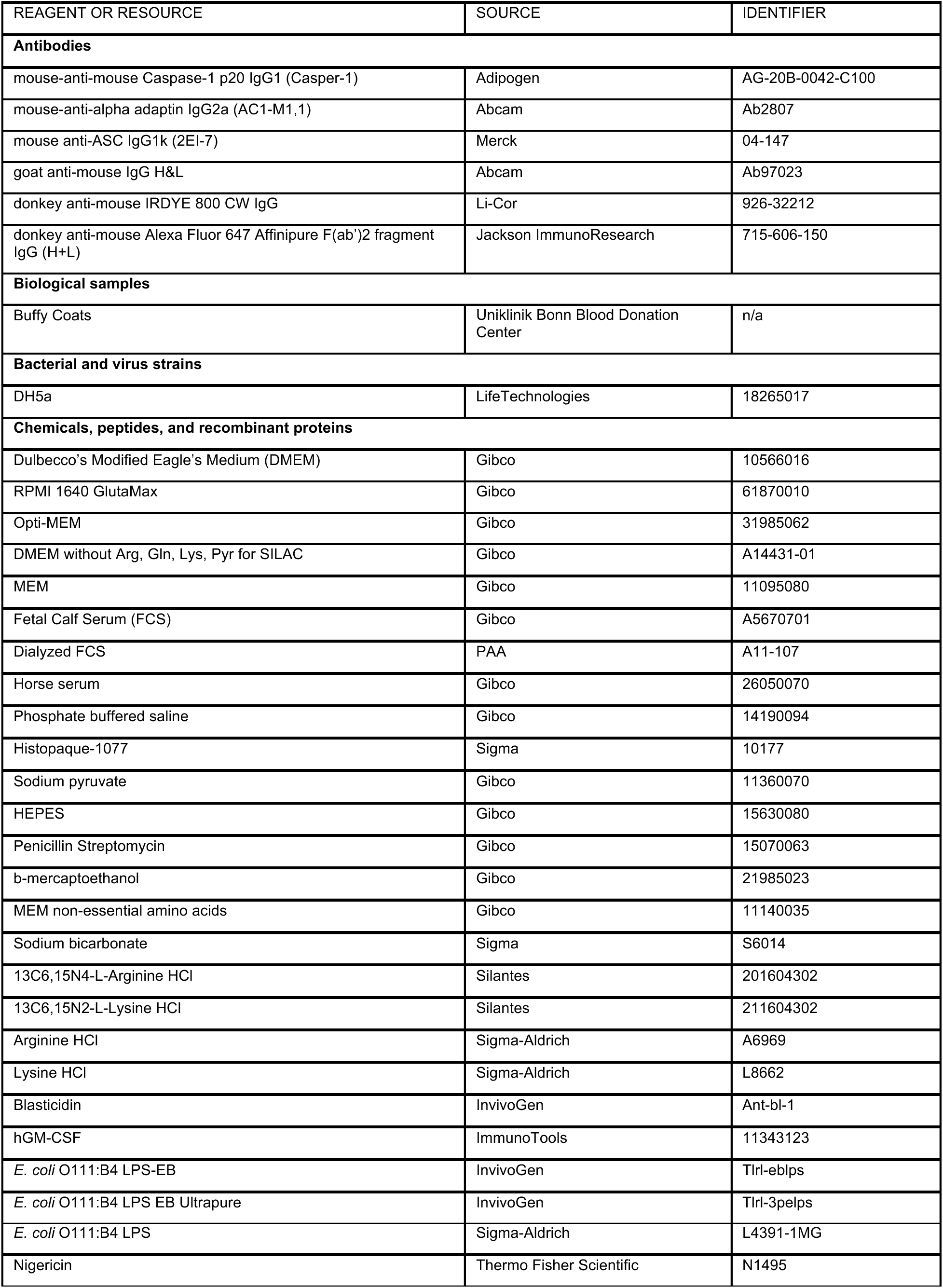
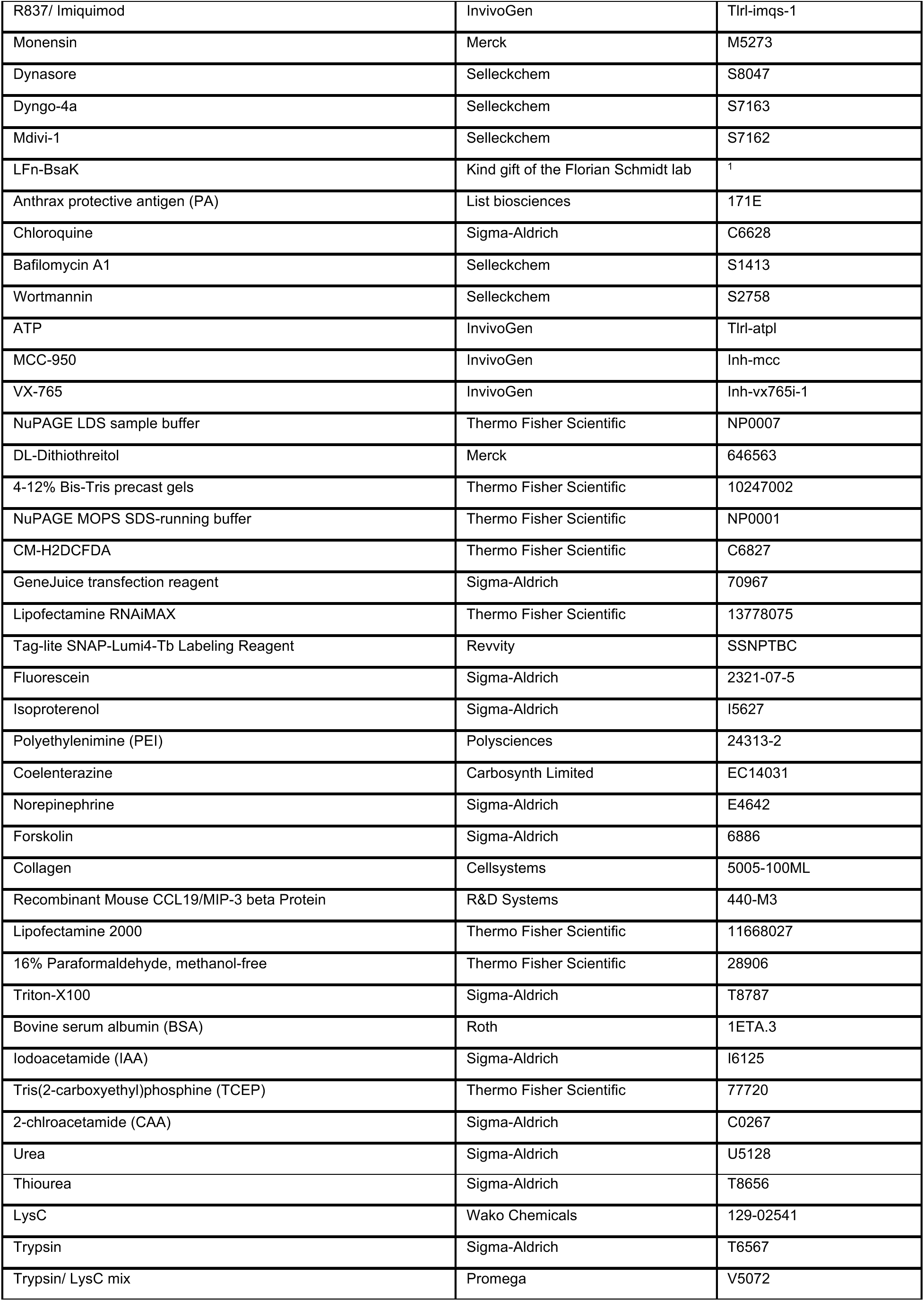
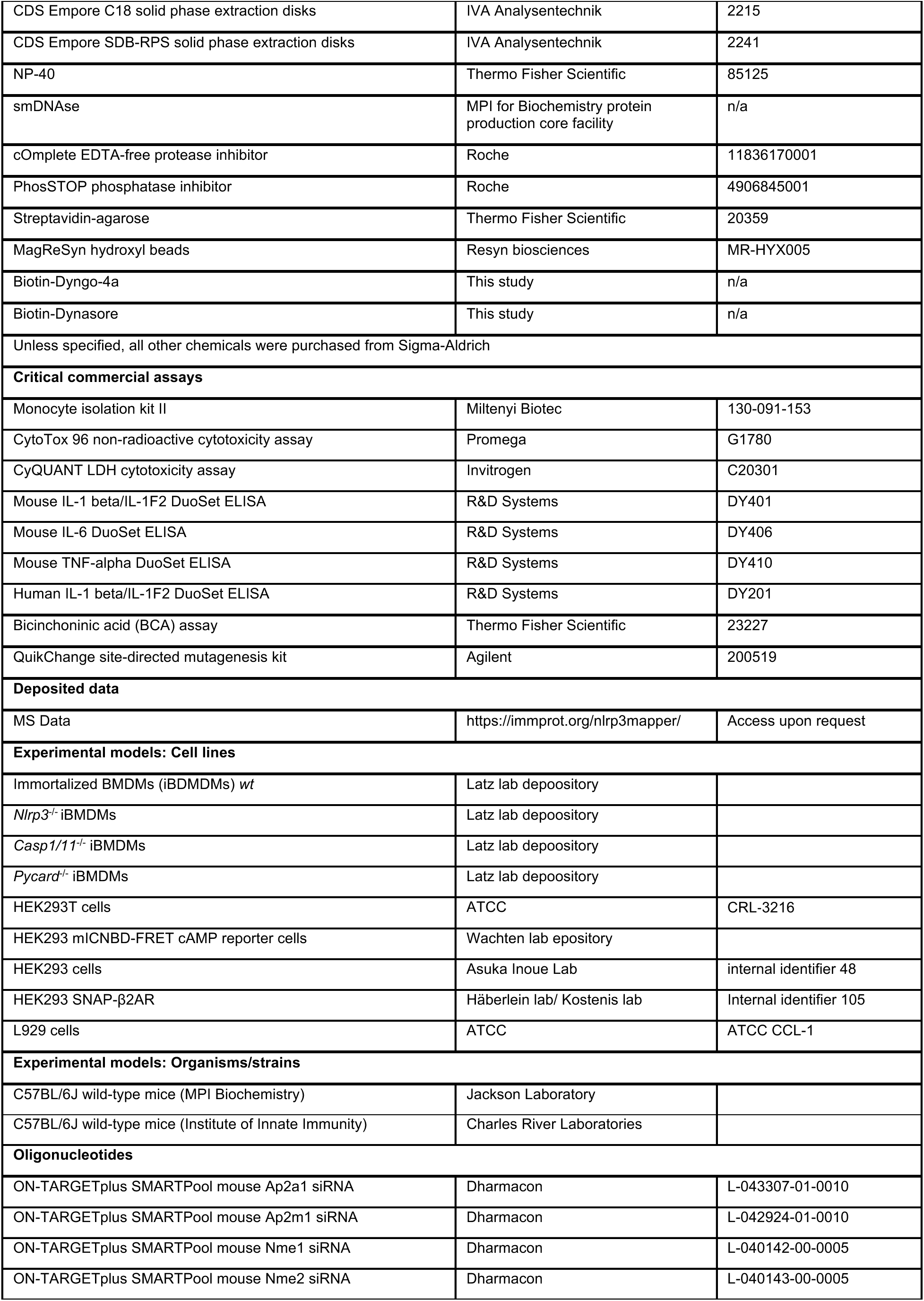
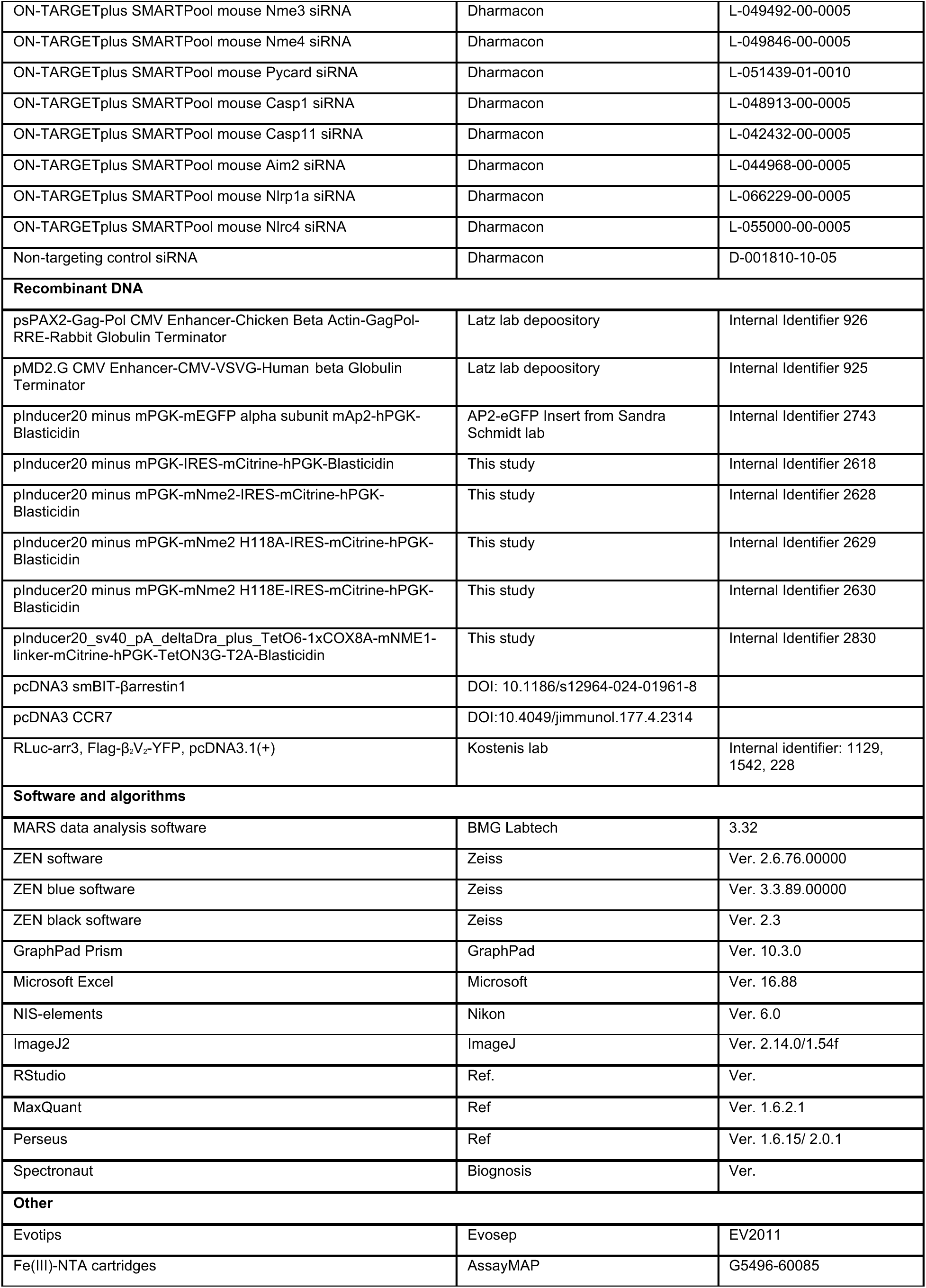
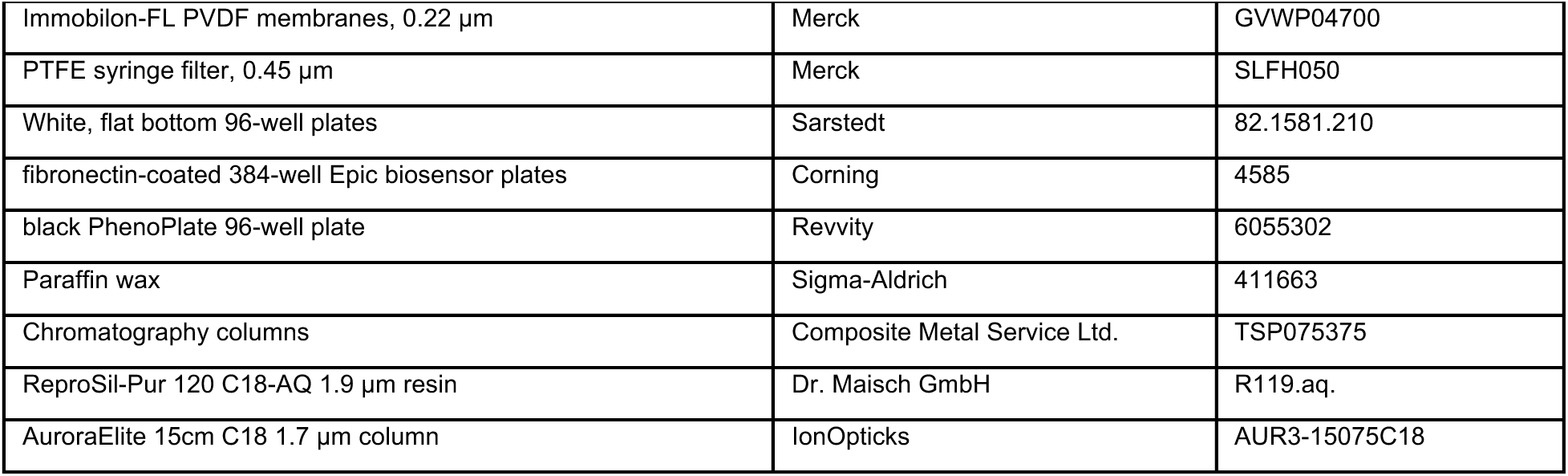
Key resource table.

## Supplementary figures

**Supplementary figure 1.**
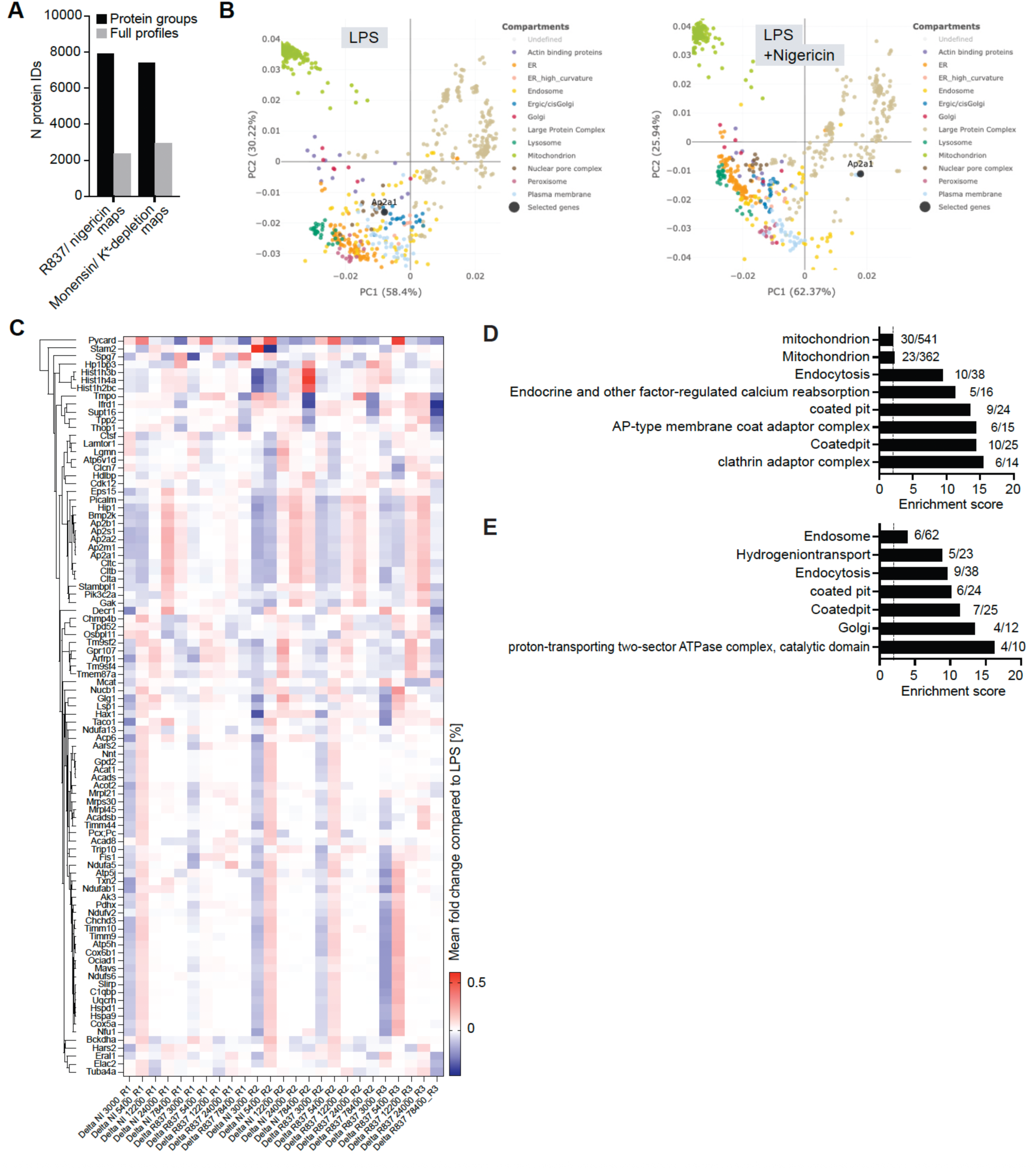
(A) Number of protein groups identified in Caspase-1^-/-^ iBMDMs before and after filtering for variance and 100% data completeness. (B) Principal component analysis of median normalized protein group L/H intensities with annotations for organellar reference proteins, exported from our webapp for LPS-treated and LPS+Nigericin treated Caspase-1-deficient macrophages. Colors indicate organellar annotations according to (*14*). (C) Heatmap of mean absolute fold change per fraction of proteins that change significantly compared to LPS control in 3 out of 5 replicates entailing both Nigericin and R837 maps. (D) Enrichment analysis for gene ontology cellular compartment (GOCC), - biological process (GOBP), -molecular function or KEGG terms and Uniprot keywords using Fisher exact test on proteins significantly (p < 0.05; FDR < 0.02, absolute fold change compared to control > 0.2) changing localization upon K^+^-depletion in ASC^-/-^ iBMDMs. (E) as in (D) for 10 µM monensin stimulation.

**Supplementary figure 2.**
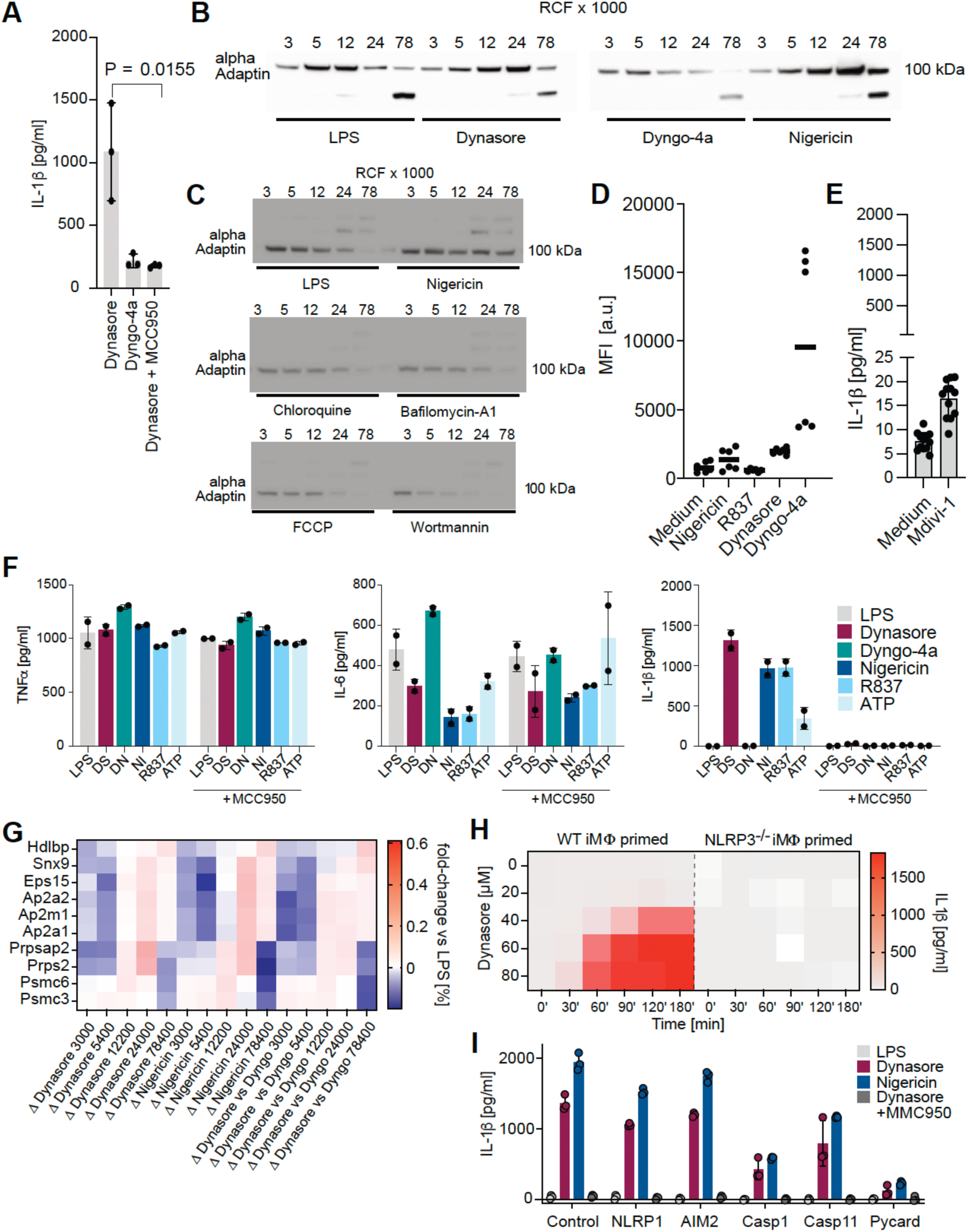
(A) IL-1β release of LPS-primed human monocyte-derived macrophages stimulated with 160 µM Dynasore ± 10 µM MCC-950 or 80 µM Dyngo-4a. Statistics indicate significance by student’s two-tailed t-test. (B) Western blot analysis of alpha-adaptin (AP2a1) in LPS-primed ASC^-/-^ iBMDMs after stimulation with 80 µM Dynasore, 40 µM Dyngo-4a or 2 µM Nigericin, using 10 µg protein per fraction per condition for individual centrifugation speeds after subcellular fractionation.(C) Western blot analysis of alpha-adaptin (AP2a1) in LPS-primed ASC^-/-^ iBMDMs after stimulation with 2 µM Nigericin, 10 µM chloroquine, 5 µM bafilomycin-4a, 10 µM FCCP or 1 µM wortmannin using 10 µg protein per fraction per condition for individual centrifugation speeds after subcellular fractionation. (D) Reactive oxygen species analysis of CM-H2DCFDA-stained ASC^-/-^ iBMDMs treated with 10 µM Nigericin, 60 µg/ml R837, 80 µM Dynasore or 40 µM Dyngo-4a for 60 min. Shown is mean fluorescent intensity. (E) IL-1β release of LPS-primed C57Bl5 BMDMs in medium or treated with a mitochondrial Dynamin DRP1-inhibitor Mdivi-1, 40 µM for 90 min. n = 3. (F) Conventional cytokine (TNFa or IL-6) and IL-1β release of LPS-primed iBMDMs treated with 80 µM Dynasore (DS), 40 µM Dyngo-4a (DN), 10 µM Nigericin (NI), 60 µg/ml R837 or 2 µM ATP or in combination with 10 µM MCC-950; n = 2. (G) Heatmap of mean fold change per fraction of proteins that change significantly comparing Dynasore to LPS, Nigericin to LPS and Dynasore to Dyngo-4a in 3 out of 3 replicates. (H) Median IL-1β release in time- and dose-titration of Dynasore treatment in *wt* or NLRP3-deficient iBMDMs, determined via ELISA. n = 3. (I) IL-1β release of LPS-primed iBMDMs which were depleted of NLRP1, AIM2, Caspase-1, Caspase-11 and ASC by siRNA stimulated with 160 µM Dynasore ± 10 µM MCC-950 or 80 µM or 2 µM Nigericin.

**Supplementary figure 3.**
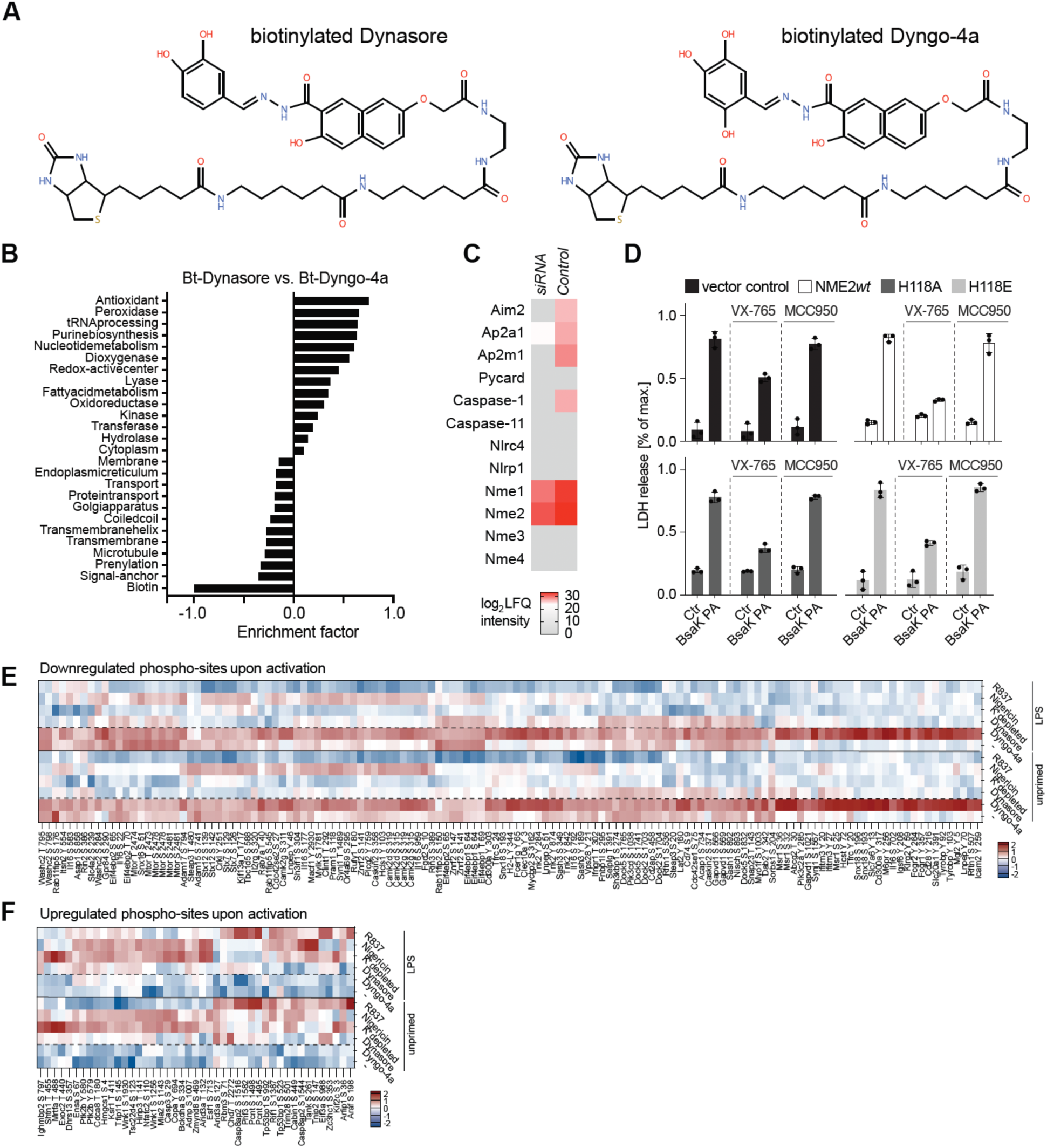
(A) Chemical structure of biotinylated Dynasore (left) and Dyngo-4a (right). (B) 1D-annotation enrichment of fold changes comparing iBMDMs treated with 100 µM bitoin-Dynasore (right) or 100 µM biotin-Dyngo-4a (left). (C) Heatmap of protein expression of knockdown targets in unprimed iBMDMs 48 h after siRNA-mediated knockdown. Shown are log2-transformed median LFQ values of n = 3; fold reduction for AP2a1 = 91.63%; NME1 = 83.65%; NME2 = 76.28%, NME3, 4 and AP2a1 were not detected after knockdown. (D) Viability (mean) of LPS-primed iBMDMs treated with the NLRC4 agonist BsaK alone, in combination with VX-765, MCC-950 or 10 µM Nigericin; n = 3. (E) Heatmap of individual regulated phosphosites according to the keywords in Fig. 3I. (F) As F, but adjacent to Fig. 3J.

**Supplementary figure 4.**
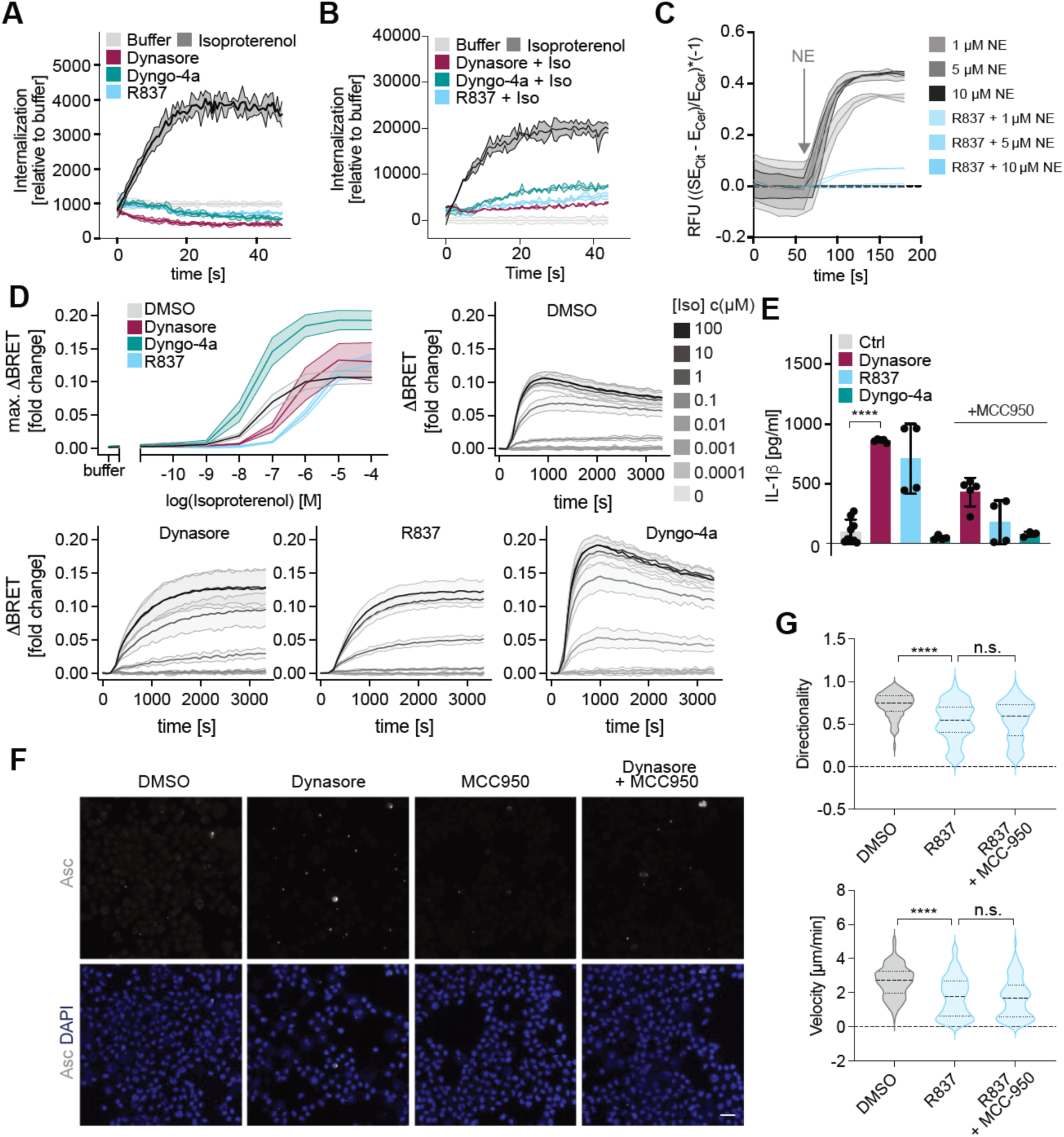
(A) Representative over-time DERET of SNAP- β_2_-AR internalization in HEK293 cells overexpressing SNAP- β_2_-AR subsequent to stimulation with 60 µM Dynasore, 30 µM Dyngo-4a, 60 µg/ml R837 or 1 µM Isoproterenol. Data are shown as the mean ± SD of one representative experiment mean ± SD of one representative experiment. (B) Representative over-time DERET of 100 nM isoproterenol induced SNAP- β_2_-AR internalization in HEK293 cells overexpressing SNAP- β_2_-AR subsequent to pre-stimulation with 60 µM Dynasore, 30 µM Dyngo-4a, 60 µg/ml R837 Data are shown as the mean. (C) Over-time cAMP production in response to norepinephrine treatment in indicated concentrations (NE). Cells were treated with DMSO or R837 (60 µg/ml) for 30 min before the experiment. (D) Maximal isoproterenol-induced recruitment of nLuc-tagged β-arrestin 2 to *wt*- β_2_-AR in HEK293 cells and individual over-time traces measured as BRET between nLuc- β arrestin 2 and a plasma-membrane mVenus-tagged kRas serving as BRET acceptor after pre-stimulation with 60 µM Dynasore, 30 µM Dyngo-4a or 60 µg/ml R837or vehicle of three independent experiments (right). (E) IL-1β release of dendritic cells stimulated with 80 µM Dynasore, 30 µg/ml R837, 10 µM Dyngo-4a or in combination with 10 µM MCC-950 for 90 min; n = 3. Statistics indicate significance by student’s two-tailed t-test (**** = P < 0.0001). (F) Immunofluorescence of ASC in BMDCs treated with 80 µM Dynasore, 10 µM MCC- 950, a combination thereof or vehicle control for 90 min. (G) Velocity in [µm/min] and directionality of dendritic cells per replicate during in-vitro 3D collagen migration assay over a timespan of 3 h after treatment with 1% DMSO, 30 µg/ml R837 or 30 µg/ml R837 + 10 µM MCC-950. Statistics indicate significance by Kruskal-Wallis test (n.s. = not significant, **** = P < 0.0001).

